# Differences in the genetic control of early egg development and reproduction between *C. elegans* and its parthenogenetic relative *D. coronatus*

**DOI:** 10.1101/171769

**Authors:** Christopher Kraus, Philipp H. Schiffer, Hiroshi Kagoshima, Hideaki Hiraki, Theresa Vogt, Michael Kroiher, Yuji Kohara, Einhard Schierenberg

**Affiliations:** Zoologisches Institut, Universität zu Köln, Cologne, NRW, Germany; National Institute of Genetics, Mishima, Japan

**Keywords:** nematode, *Caenorhabditis*, parthenogenesis, embryogenesis, polarity, chromosome, genome, transcriptome, hybridization

## Abstract

**Background:** The free-living nematode *Diploscapter coronatus* is the closest known relative of *C. elegans* with parthenogenetic reproduction. It shows several developmental idiosyncracies, for example concerning the control of meiosis and embryonic axis formation [1]. Our recent genome analysis [2] provided some support for the view that *D. coronatus* is a product of interspecies hybridization. Here we present additional data towards this assumption. Based on genomic and transcriptomic data we try to better understand the molecular basis of developmental idiosyncrasies in this species in an evolutionary context by comparison with selected other nematodes.

**Results:** In a genomic comparison between *D. coronatus*, *C. elegans*, other representatives of the genus *Caenorhabditis* and the more distantly related *Pristionchus pacificus* and *Panagrellus redivivus*, certain genes required for normal embryogenesis in *C. elegans* were found to be restricted to the genus *Caenorhabditis.* The mRNA content of early *D. coronatus* embryos was sequenced and compared with similar stages in *C. elegans* and *Ascaris suum.* We identified 350 gene families transcribed in the early embryo of *D. coronatus* but not in the other two nematodes. Looking at individual genes transcribed early in *D. coronatus* but not in *C. elegans* and *A. suum* we found that orthologs of most of these are present in the genomes of the latter species as well, suggesting heterochronic shifts with respect to expression behavior. Considerable divergence between alleles lends further support to the view that *D. coronatus* may be the result of an interspecies hybridization. Expression analysis of early acting single copy genes yield no indication for silencing of one parental genome.

**Conclusions:** Our comparative cellular and molecular studies support the view that the genus *Caenorhabditis* differs considerably from the other studied nematodes in its control of development and reproduction. The easy-to-culture parthenogenetic *D. coronatus*, with its high quality draft genome and only a single chromosome when haploid, offers many new starting points on the cellular, molecular, and genomic level to better understand alternative routes of nematode development and reproduction.

## Background

Nematodes can follow different modes of reproduction including parthenogenesis. This reproductive mode is a deviation of an original bisexual situation and has been established several times independently within different metazoan taxa. It can arise in several ways including spontaneous mutation, interspecies hybridization or infection with microorganisms and may go along with regular meiosis followed by fusion of gametes, or complete or partial suppression of meiosis[3]. However, no parthenogens have been found in the genus *Caenorhabditis*, despite the rapidly rising number of described species (>30) while at the same time a shift from gonochoristic to hermaphroditic reproduction took place in this taxon several times independently [4,5]. Development of the model *Caenorhabditis elegans* has been extensively studied. Although comparative studies in other nematodes revealed considerable variations on the cellular level (for review, see [6]) it seemed self-evident that gene cascades controlling development are conserved across the phylum. However, analysis on the levels of genome and transcriptome suggested major changes in the logic of cell specification and the action of Developmental System Drift [7] even between nematodes from neighboring clades [8,9]. While *C. elegans* can obviously not serve as a general model for nematode development it has remained unclear how fast the genetic control of development has changed during evolution in the long-branched roundworms.

Therefore, we here analyze molecular and cellular aspects of early development and reproduction in the parthenogenetic species *Diploscapter coronatus*, which has just half the body size of *C. elegans* and whose genome we described recently [2]. *D. coronatus* is a member of the Protorhabditis group, which not only belongs to the same clade as the genus *Caenorhabditis* but is the immediate sister taxon of it [10]. We previously described some idiosyncrasies in early development of *D. coronatus* using microscopic approaches [1,11].

In the androdioecious hermaphrodite *C. elegans* oocytes arrest in meiotic prophase and are released sequentially, this way delivering a continuous supply of maturing oocytes {McCarter:1999bu, [12]. The generation of somatic founder cells via asymmetric germline divisions in *D. coronatus* takes place in the same way as in *C. elegans* despite the absence of sperm-induced polarization prior to first cleavage. In *D. coronatus*, only one polar body is generated during a truncated meiosis explaining the diploid status without fertilization. This suggests differences in the molecular machinery initiating axis polarity.

The control of oocyte maturation in *C. elegans* requires signaling from the sperm via Major Sperm Protein (MSP) [13]. We found earlier that MSP genes are present in parthenogenetic nematodes, including *D. coronatus*, however, MSP protein could not be detected there [14].

Screening the gene and protein sets of *D. coronatus* for regulators of important developmental processes in *C. elegans* we make comparisons with other members of the genus *Caenorhabditis*, as well as two more distantly related nematodes with gonochoristic and hermaphroditic reproduction. Particularly, we search for peculiarities that can be related to the development of oocyte and early embryo in the context of parthenogenetic reproduction in *D. coronatus.*

In a second approach, we compare the transcriptome of early embryonic stages in *D. coronatus* with the known complement of genes expressed in corresponding stages of *C. elegans* [15] and *Ascaris* [16]. In particular, we are interested to explore to what extent the expression of certain genes in *D. coronatus* can be correlated to its early developmental idiosyncrasies.

With our earlier finding in mind that the genome of *D. coronatus* shows a high degree of heterozygosity [2], we looked for further evidence that parthenogenesis in this species may be the result of interspecies hybridization which is considered a major route to this mode of reproduction in invertebrates and possibly the only one in vertebrates [17].

## Methods

### Nematode culture and strains

Strains were cultured on agar plates with the uracil-requiring OP50 strain of *E. coli* as a food source, essentially as described by Brenner[18], except that, to reduce contamination with other bacteria, we used minimal medium plates [11]. *Diploscapter coronatus* (PDL0010) was kindly provided by Paul De Ley, Dept. Nematology, University of California, Riverside.

To measure brood size, 17 juveniles of *D. coronatus* were isolated and grown individually as described above. When starting to lay eggs, animals were transferred to new culture plates every two days until they died and the total of hatched larvae was counted.

### Microscopical analysis and 3-D reconstructions

Development was studied with Nomarski optics using a 100x objective. 1-cell stage embryos were collected from agar plates with a drawn-out Pasteur pipette or after dissection of gravid adults. Specimens were placed on microscope slides carrying a thin agar layer as a mechanical buffer and covered with a coverslip sealed on the edges with petroleum jelly. Development was recorded using a 4-D microscope with 15-25 optical sections/embryo and 15-60 sec time intervals between scans [19]. Cell behavior was traced with help of the Simi Biocell software (Simi Reality Motion Systems GmbH, Unterschleißheim, Germany). Nuclei were counted in optical sections of DAPI-stained isolated gonads.

### *D. coronatus* ITS, SSU, LSU rDNA analysis

For each *D. coronatus* rDNA gene two individuals were picked and lysed. Using single-worm PCR[20] we cloned sequences from each rDNA gene and individual into separate pBluescript KS cloning-vectors. For amplification of the ribosomal small subunit (SSU) we used primers described in [21] and [22], for the ribosomal large subunit (LSU) primers from [23] and for the ribosomal internal transcribed spacer (ITS) from [24]. For phylogenetic analysis we used Mr. Bayes (Ronquist and Huelsenbeck, 2003; version 3.1.2) and RAxML (version 7.2.8) [25] with standard parameters and 100 bootstraps. Resulting trees were collapsed after first node.

### OrthoMCL clustering and identification of the presence and absence of orthologs

To reliably compare orthologues we used the OrthoMCL clustering pipeline (version 2.0.9)[26] including proteomes of five *Caenorhabditis* species (*C. angaria*, *C. briggsae*, *C. elegans*, *C. japonica*, *C. tropicalis*; [27-29]; http://www.ebi.ac.uk/ena/data/view/GCA_000186765.1). *Diploscapter coronatus*, *Pristionchus pacificus* [30], *Panagrellus redivivus* [31] and *Ascaris suum* [32],[16]. The absence of genes in the *D. coronatus* genome which are present in the genomes of *C. elegans*, *C. briggsae and C. remanei* was confirmed by reciprocal BLAST search.

### Gene ontology (GO) term analysis

Fisher’s exact test for gene ontology (GO) terms of *D. coronatus*-*specific* clusters and singletons (proteins comprising a species-specific single variant) during early embryogenesis was applied to identify significantly over-represented GO terms [33,34] (FDR < 0.05; p < 0.001).

### Phylogenetic classification and analysis

We here refer to the phylogeny of Holterman[35], dividing nematodes into 12 different clades. Following De Ley and Blaxter [36] we distinguish more basal Enoplea (clades 1 and 2) from more derived Chromadorea (clades 3-12). While *C. elegans*, *D. coronatus* and *Pristionchus pacificus* are members of clade 9, other species mentioned in this paper belong to clade 12 (*Meloidogyne* spp.), clade 11 (*Acrobeloides nanus*), clade 10 (*Panagrellus redivivus; Panagrolaimus* spp.), clade 8 (*Ascaris suum*), clade 6 (*Plectus sambesii*) and clade 2 (*Romanomermis culicivorax*).

In order to visualize the structural differences between the *C. elegans* and the *D. coronatus* LET-99 homologs multiple alignments were performed using the program Clustal OMEGA[37]. Outgroup proteins including an N-terminal DEP-domain were retrieved from Pfam database[38]. For the phylogenetic analysis, the best amino acid (AA) substitution matrices were identified using the program Prottest3 under the conditions of invariant sites and gamma optimization. Best substitution matrices were identified under the condition of the Bayesian information criterion and the Akaike information criterion[39]. Phylogenetic trees were constructed using RAxML with gamma value optimization and the substitution matrices suggested by Prottest3. Each tree was bootstrapped 100 times.

### RNA extraction and RNA sequencing of selected embryonic stages

For RNA sequencing we collected under the dissecting scope four batches (= 4 independent biological replicates) of approximately 100 eggs each, consisting of 1-8 cell stage embryos. These were placed into 25 μl H_2_O, shock-frozen in liquid nitrogen and immediately stored at −80°C to avoid RNA degeneration. RNA of each sample was extracted by a slightly modified version of an established protocol [40]. Instead of using 4M Guadiniumthiocyanate (GU) buffer, we used 6M GU buffer. By adding 175 μl 6M GU buffer and using a homogenizer (Ultra-Turrax, IKA Werke GmbH) it was possible to lyse the samples under chaotropic conditions. The extracted amount of total RNA was dissolved in 2μl RNAse free water and used for RNA amplification using the “Message AMP II” kit (AM1751; Life Technologies Inc.) following the protocol of Hashimshony et al., (2012). This allowed linear amplification (in contrast to exponential amplification methods such as PCR) of the total RNA content, hence significantly decreasing the amplification bias. TruSeq library construction (TruSeq preparation kit version 2; Illumina Inc.) and RNA sequencing was performed on Illumina HiSeq and MiSeq platforms at the local sequencing facility (CCG Cologne). Retrieved paired-end reads ranged from approximately 8,500,000 to 31,000,000; depending on the sequencing platform.

### Post-sequencing analysis

Illumina paired-end reads were retrieved in four independent sequencing assays. Illumina adapters and indexes were removed using the program Trimmomatic [41] and 5′ and 3′-prime error-prone reads were removed using the program sickle (github.com/najoshi/sickle). Trimmed reads were used to generate a transcriptome using the *de*-*novo* assembler Trinity[42]. To identify even scarce transcripts all four libraries were combined, this way a transcriptome with a maximum number of transcripts and the highest median was obtained. To screen for and eliminate bacterial contamination, assembled transcripts were mapped back to the *D. coronatus* EST library and transcriptome using bowtie2[43]. For comparison with similar early embryonic stages of *C. elegans* and *A. suum* the raw data were taken from [15] and [44]. In the case of *C. elegans* sequences showing an average TPM (transcripts per millions) value of >5 were counted as being expressed. In *A. suum* significant expression differences between the 1- to 4-cell stages on the one hand and subsequent stages on the other [44] allowed an estimation of the early stage-specific transcriptome. The transcriptome of *D. coronatus* was translated into protein sequences using the program Transdecoder (Haas et al., 2013) with a minimum AA length of 49 residues. Transcripts for which information was only available for the 3’-UTR (untranslated region) or which where shorter than 49 residues were aligned to an EST library to generate extended gene models. Resulting extended contigs were translated into AA sequences following the same procedure as described above. For *C. elegans* and *A. suum* proteins corresponding to the early transcriptome were downloaded from wormbase.org. The retrieved protein sequences were used for orthologous clustering using OrthoMCL.

### Identification of “allelic” gene variants

We determined the intron-exon structure and the positions on the genome for all of the predicted genes. By aligning and clustering all EST libraries making use of CD-HIT[45] at a threshold of 90% identity, we identified ESTs belonging to the same gene. We mapped clustered ESTs against the genome using BLAT (Kent, 2002) this way confirming the exact position of each EST cluster on the genome. EST clusters mapping to open reading frames (ORFs) of genes were translated into AA sequences. Corresponding proteins were cross-compared by an all-vs.-all blast [46] approach (at a threshold of 98% identity). We sampled pairwise occurring genes with an amino-acid identity of >98%. These were considered as different copies of the same gene under the condition that they were positioned on different contigs. In the following we call these pairwise occurring genes with high AA identity “alleles”. Taking into account the relative position on the genome, we deduced their numbers by counting positions and contigs.

### Identification of *C. elegans* genes in *D. coronatus* with two “alleles” at different loci

Orthologous clusters consisting of only two proteins were extracted from our OrthoMCL clustering. Respective protein sequences were pairwise aligned via clustal OMEGA multiple alignment algorithms. AA identity was calculated by custom Perl scripts. Sequences which are known to exist at two different loci in the *C. elegans* genome and still have an identity of 95% were considered to be genes existing in two distinct “alleles”.

### Single-copy gene analysis

Single-copy genes present in nematodes of various clades as well as in *Drosophila melanogaster* and *Saccharomyces cerevisiae* were selected based on [47]. *D. coronatus* and *C. remanei* orthologs were identified by OrthoMCL. For *C. remanei* candidates were retrieved from wormbase.org using the *C. elegans* orthologs described in [47]. Known *C. elegans* orthologs were identified in the *D. coronatus* genome with NCBI Blast. The identity of a *D. coronatus* ortholog was confirmed by OrthoMCL clustering and by using the predicted genes of *D. coronatus* as a query to search for the original ortholog in the *C. elegans* genome (best-reciprocal-blast-hit approach).

We compared orthologs by pairwise alignments of *D. coronatus* “alleles” against each other and the *C. elegans* alleles against the *C. remanei* sequence in clustalW [48].

Orthologs of single-copy genes were scanned for conserved protein domains with InterProScan [49]. We counted the number of synonymous and non-synonymous mutations within the respective single-copy genes by pairwise alignment of the sequences on the nucleotide level taking into account the appropriate reading frame and by using the KaKs Calculator (version 1.2) with standard parameters (http://evolution.genomics.org.cn/software.htm; [50]). For a statistics of average nucleotide exchange rates, the one-tailed Welch t-test for non-equal variances was applied. We tested specifically the null hypothesis, i.e. whether the fraction of non-synonymous mutations is equal or greater than the fraction of synonymous mutations. The null hypothesis was rejected at a significance level of α=0.01.

### Identification of expressed *D. coronatus* “alleles” during early embryogenesis

*D. coronatus* transcripts expressed during early embryogenesis were mapped back to the *D. coronatus* expressed sequence tag (EST) library using the program Bowtie2. Transcripts mapping to two different genomic contigs were considered as “alleles” (see above). ESTs were clustered using the program CD-HIT with standard parameters for >94% identity. Mapped reads were used to identify polymorphisms (in particular single nucleotide polymorphisms; SNPs) by using the programs SAMtools and Bcftools [51] with a minimum sequencing coverage of 10-fold per site. The genotype quality (GQ, [52]) of each retrieved SNP was represented by a maximum likelihood for wrong SNP calls of <10^-3^. Variations which did not meet these criteria were considered as random variants probably due to sequencing errors. Halplotypes for each variant were inferred by usage of the “vcfgeno2halpo” command of the Vcflib suite (https://github.com/vcflib/vcflib) for a window size of 500 bp. Transcripts with >99% nucleotide identity were defined as indistinguishable and excluded from the analysis.

## Results

### Multi-species orthologous clustering

We compared our *D. coronatus* data with other nematodes, including *C. elegans*, to better understand the molecular basis of developmental peculiarities in this species (Table 1). The *D. coronatus* draft genome contains more than 34,000 protein predictions and we used these to screen for conserved and species-specific genes. In order to identify robust orthologous clusters in comparison to several other species selected for their phylogenetic position (Table 1, Fig. 1) we used OrthoMCL. In total, we found over 8,000 orthologous clusters shared between *P. redivivus* (clade 10) and *A. suum* (clade 8). About 80 % of these are present in all seven clade-9 species considered here as well, suggesting a core set of shared protein families. However, the majority of the nearly 20,000 clusters are not shared with *P. redivivus* and *A. suum* (Fig. 1).

**Table 1.** Species and proteomes used in this study.

| species | clade<sup>a</sup> | number of proteins |  |
| --- | --- | --- | --- |
|  |  | whole genome | early embryogenesis |
| *C. elegans* | 9 | 22,855 | 8,817 |
| *C. angaria* | 9 | 28,371 | - |
| *C. japonica* | 9 | 30,083 | - |
| *C. tropicalis* | 9 | 24,532 | - |
| *D. coronatus* | 9 | 31,693 | 4,610 |
| *P. pacificus* | 9 | 23,750 | - |
| *A. suum* | 8 | 17,555 | 3,093 |
| *P. redivivus* | 10 | 26,372 | - |

<sup>a</sup>, for phylogenetic classification, see Methods

**Fig. 1:**
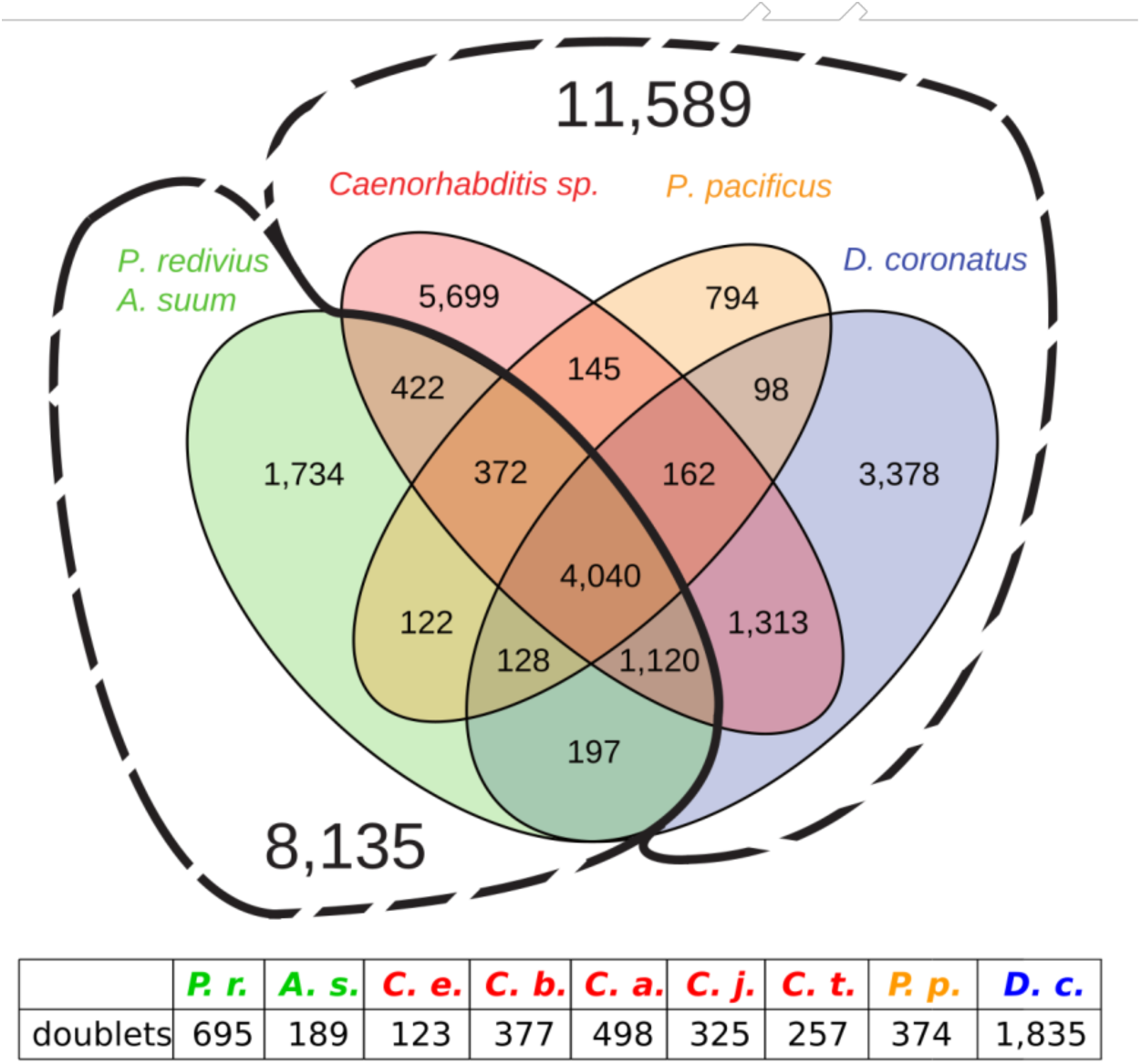
a, Distribution of shared and specific orthologous clusters for representatives of *Diploscapter coronatus* (blue), *Pristionchus pacificus* (orange) and the genus *Caenorhabditis* (red), as well as for the outgroups *Panagrellus redivivus* (clade 10) and *Ascaris suum* (clade 8; both in green). Insert: Number of species-specific proteins present in two distinct “alleles” (“doublets”).

By analyzing five *Caenorhabditis* species and in addition at *D. coronatus*, *P. pacificus and Panagrellus redivivus*, we found that over 5,000 orthologous clusters, or nearly 50% of all clade 9-restricted clusters were specific to the genus *Caenorhabditis.* This suggests that during *Caenorhabditis* evolution a considerable number of genes must have newly arisen in the lineage leading to this taxon.

### Absence of genes and development in *D. coronatus*

Using our ortholog clustering (Fig. 1) we investigated which genes known to be crucial in development of *C. elegans* are restricted to the genus *Caenorhabditis.* We found an absence of developmental regulators for a variety of biological processes (Fig. 2) and decided to focus on oogenesis and early embryogenesis where we had observed phenotypical idiosyncrasies in *D. coronatus.*

**Fig. 2:**
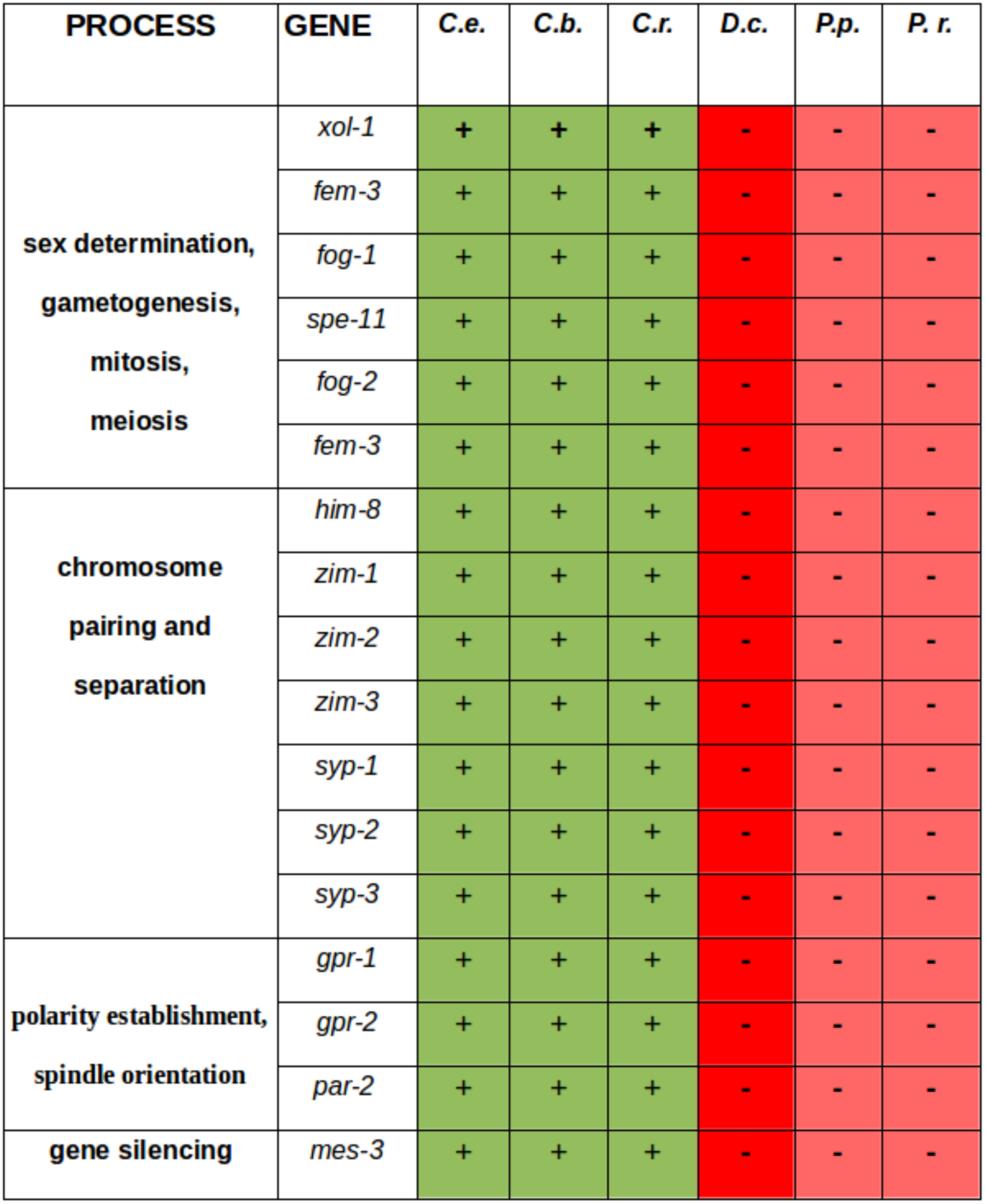
Orthologs of genes with essential functions in different processes in *C. elegans* not found in *D. coronatus* and two other species outside the genus *Caenorhabditis*. Green, genes found; red, genes not found. *C.e*, *C. elegans*; *C.b*, *C. briggsae; C.r.*, *C. remanei; D.c*, *D. coronatus*; *P.p.*, *P. pacificus*; *P.r*.; *P. redivivus*.

### Oocyte development and modified meiosis

In *C. elegans* the development of germ cells from mitotic oogonia to mature oocytes is arrested in prophase of meiosis I and their subsequent activation by sperm is an elaborate process [53]. We found that each of the two gonadal arms of the mature *D. coronatus* adult is about 5x smaller than in *C. elegans* and contains only 30-100 germ cell nuclei (Fig. 3a,b; n=20) in contrast to the latter where about one thousand are generated [54]. Under our laboratory conditions, individual *Diploscapter* females produced less than one third the number of eggs found in *C. elegans* (on average 80; n=17). Different to the latter, the size of developing germ cells in *D. coronatus* increases only marginally except for the most mature one (“-1 oocyte”) which is much larger and densely filled with yolk granules (Fig. 3a; in older adults also the −2 oocyte starts to grow). We wondered whether the same phases of oocyte differentiation as in *C. elegans* can be found in *D. coronatus.*

**Fig. 3:**
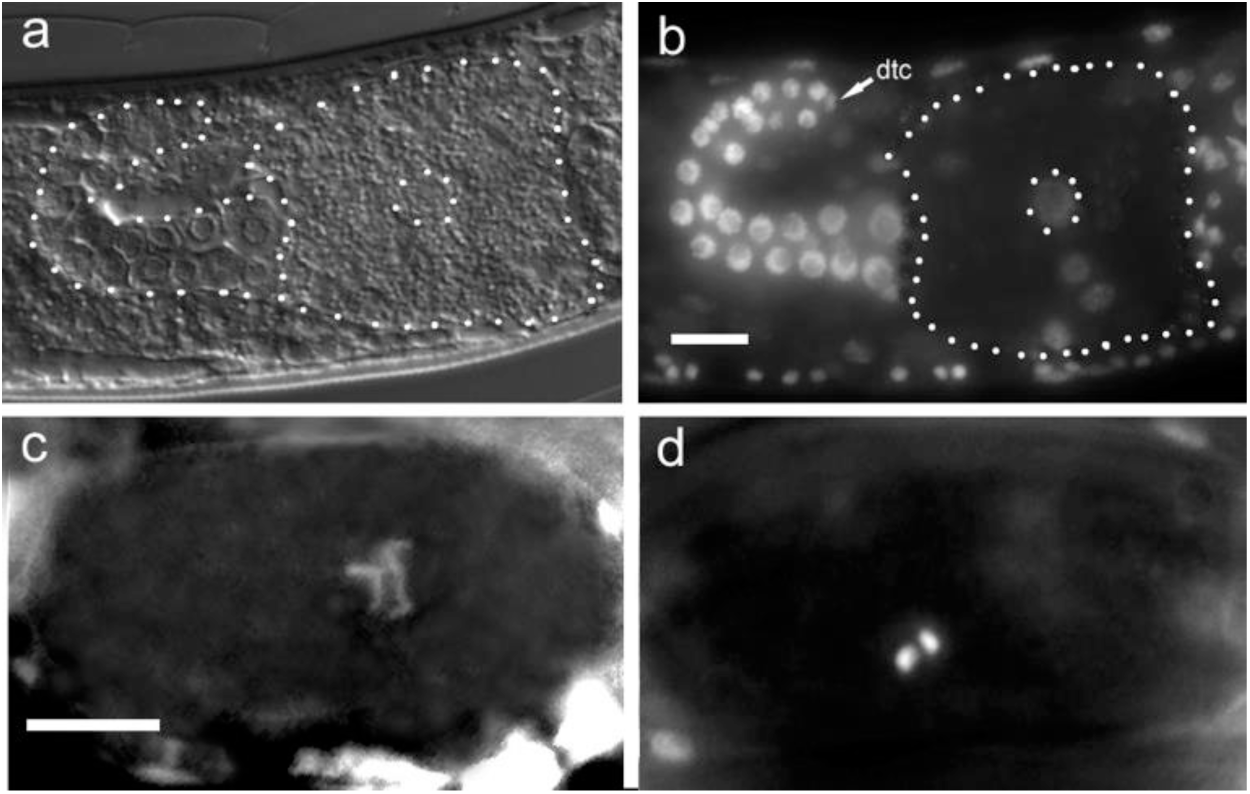
Gonad and chromosomes in *D. coronatus.* a, outline of one gonadal arm including one large oocyte; b, gonadal arm with DAPI-marked germ cell nuclei; c, d, extended and condensed chromosomes in prophase of meiosis I.

The analysis of DAPI-stained germline nuclei indicates that this is not the case. In the adult *D. coronatus* ovary essentially all germ cell nuclei appear to be in premeiotic interphase (Fig. 3b) except for rarely observed mitoses in the distal-most region. Condensed meiotic chromosomes were only found in oval-shaped 1-cell stages in the uterus surrounded by an eggshell (Fig. 3c, d). Thus, in contrast to *C. elegans*, in *D. coronatus* individual germ cells seem to enter meiosis late and one-by-one without prophase arrest.

*D. coronatus* contains only two chromosomes in the diploid status (2n=2; Fig. 5c, d) [55], whereas in *C. elegans* 2n equals 12 chromosomes [56]. In accordance with Hechlers report we never observed paired meiotic chromosomes.

Screening the *D. coronatus* genome for genes essential for germ cell development or sex-specific cell differentiation in *C. elegans*, we found orthologs for several of such genes missing. However, their absence cannot serve as a straight forward explanation for special features of the parthenogenetic *D. coronatus* as they were not detected in *Pristionchus* and *Panagrellus* as well (for phylogeny, see Methods), while present in all three considered *Caenorhabditis* species (Fig. 2). At least, our data indicate that the control of central developmental processes differs between members of the genus *Caenorhabditis* and representatives of neighboring clades and even within the same clade.

#### Polarity establishment and early embryogenesis

Gonochoristic and hermaphroditic reproduction depends on sperm, which contributes the centriole, essential for mitotic spindle formation, and initiates embryonic polarity as a prerequisite for subsequent asymmetric cell division and soma/germline separation in *C. elegans* [57,58].

The proper positioning of the first cleavage spindle in *C. elegans* and subsequent movement of one aster towards the posterior pole of the zygote preceding its asymmetric division requires the presence of LET-99. The C-terminal domain of LET-99 appears to be important for its functionality (Fig.4) as nonsense mutations lead to strong phenotypes [59]. LET-99 is also the main regulator for the localization of LIN-5 and GPR-1/GPR-2 [60] which act together to generate the necessary spindle pulling force [61,62]. In the genome of *D. coronatus* we could not find orthologs of *gpr*-*1/2* (Fig. 2).

**Fig. 4:**
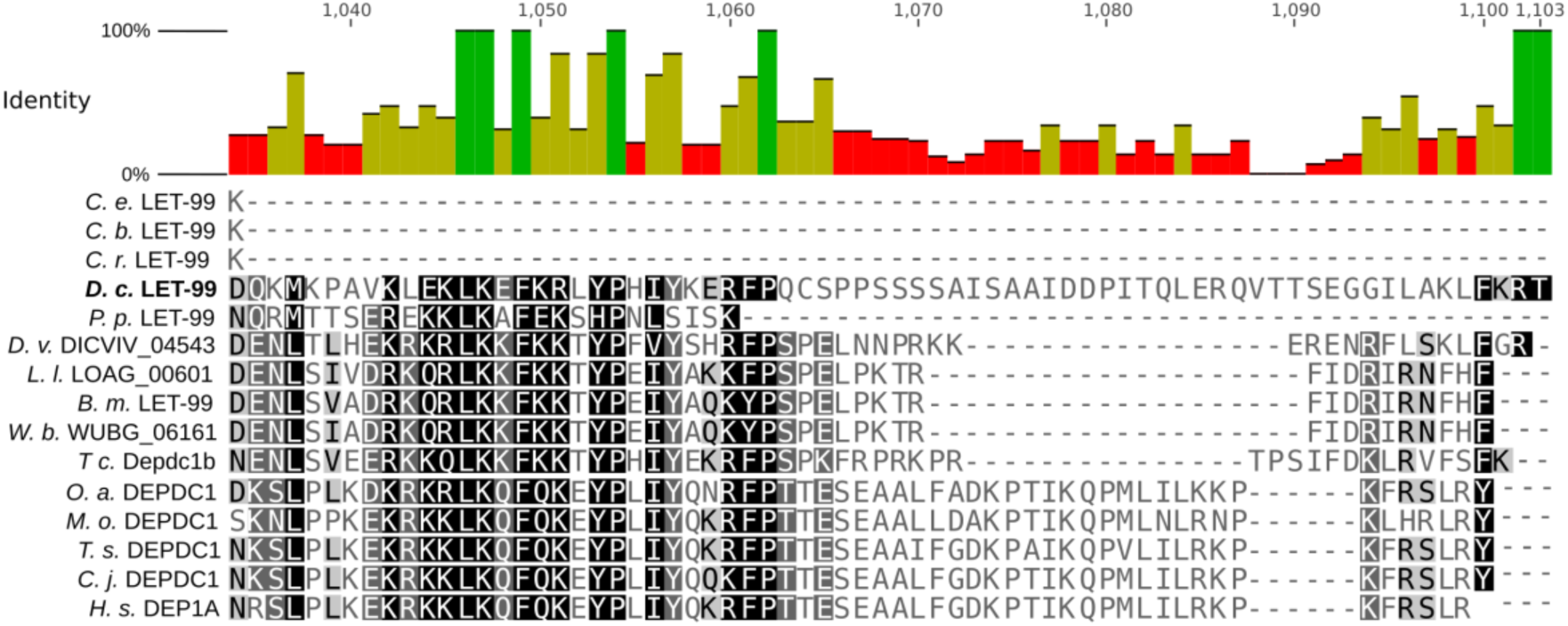
Sequence alignment of the C-terminal region of the *D. coronatus* LET-99 domain in comparison to *Caenorhabditis*, other nematodes and the homologous mammalian DEPDC1 protein. The C-terminal region is absent in *Caenorhabditis* species (*C. e.*, *C. elegans*; *C. b.*, *C. briggsae*; *C. r.*, *C. remanei*) but present in the other tested nematodes (*D. c*, *D. coronatus; P. p.*, *Pristionchus pacificus*; *D. v.*, *Dictyocaulus viviparus*; *L. l.*, *Loa loa*; *B. m.*, *Brugia malayi*; *W. b.*, *Wucheria brancrofti*; *T. c.*, *Toxocara canis*) and vertebrates (*O. a.*, *Ornithorhynchus anatinus*; *M. o.*, *Microtus ochrogaster*; *T. s.*, *Tarsius syrichta*; *C. j.*, *Callithrix jacchus*; *H. s.*, *Homo sapiens*).

We identified a LET-99 homolog in *D. coronatus* (DcLET-99). Alignment of the protein sequence with that of other species revealed, however, that the last 70 AA of the C-terminal region are absent in the genus *Caenorhabditis*. The *Diploscapter* ortholog thus has more similarity with the protein in other nematodes, like the ones included in our study (Fig. 4). This finding may reflect a nonequivalent function of this protein in *C. elegans* and *D. coronatus*.

With these findings in mind, we compared the establishment of asymmetry between *D. coronatus* and *C. elegans.* Characteristic for *C. elegans* is the migration of the two pronuclei to the center of the fertilized egg cell and their subsequent fusion (Fig. 5a-c). Consequently, the posterior aster together with the future P_1_ chromosome set is quickly translocated to the posterior while the anterior aster remains at its original position (Fig. 5c, d). This results in an asymmetric division of the zygote into a larger somatic AB and a smaller P_1_ germline cell (Fig 5e, f). Subsequently, AB divides with transverse and P_1_ with longitudinal spindle orientation (Fig. 5g) resulting in a rhomboid 4-cell stage (Fig. 5h). With the division of P_2_ a reversal of cleavage polarity (PR) takes place in the germline such that P_3_ occupies a more anterior position relative to its somatic sister C (Fig. 5i) [63].

**Fig. 5:**
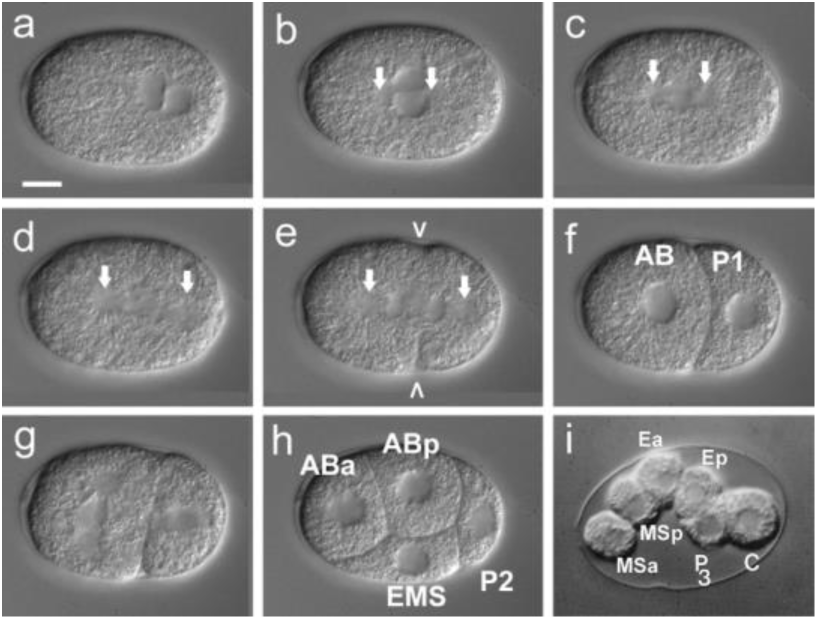
Early embryogenesis of *C. elegans.* For details, see text. Arrows, centriolar regions; arrowheads, cleavage furrow; i, for better visualization of polarity reversal in the germline AB cells had been removed through a laser-induced hole in the eggshell. Scale bar, 10 μm; orientation, anterior, left.

In the parthenogenetic *Diploscapter* only one pronucleus is generated during meiosis (Fig. 6a, b; Lahl et al., 2006). In contrast to *C. elegans*, a temporary constriction forms at the anterior pole and the maternal pronucleus occupies a slightly eccentric position (Fig. 6b-d). The zygote divides with no shift of the posterior aster (Fig.6c-e) while the constriction regresses. This way, a larger AB and a smaller P_1_ cell are formed (Fig. 6e-f). Subsequently, AB divides with longitudinal spindle orientation like P_1_ (Fig. 6g, h) and a PR in P_2_ is absent (Fig. 6i). The absence of *gpr*-*1* and *gpr*-*2* plus the considerably diverged *let*-*99* could give an explanation for the different ways of how the asymmetric division of the zygote is achieved in the two species.

**Fig. 6:**
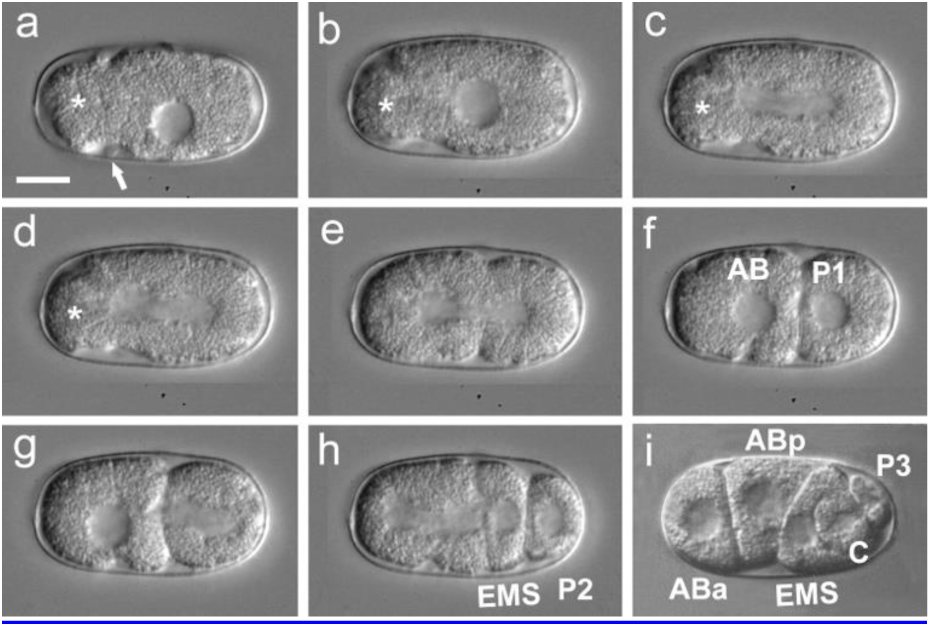
Early embryogenesis of *D. coronatus.* For details, see text. Arrow, single polar body; asterisk, separated anterior cytoplasm. Scale bar, 10 μm. Orientation, anterior, left.

As *gpr*-*1* and *gpr*-*2* are absent in *P. pacificus* and *P. redivivus*, too (Fig. 2), we studied formation of asymmetry in 1-cell embryos there and found it to be similar to *D. coronatus* while *C. briggsae* and *C. remanei* behave like *C. elegans.* This can also be deduced from the video clips accompanying Brauchle et al. (2009). In addition, we analyzed one representative each of Cephalobidae (*A. nanus*) and Plectidae (*Plectus sambesii*). With respect to spindle movement they behave similar to *D. coronatus* (data not shown). These findings indicate that the genus *Caenorhabditis* has developed a special way of how to accomplish the first asymmetric cleavage.

### Spindle orientation and germline polarity

As the same a-p spindle orientation in the AB cell of *D. coronatus* was also found in a *par*-*3* mutant of *C. elegans* [64] we screened the genome of *D. coronatus* for the presence of *par* genes. We found an ortholog of *par-3* but not of *par*-*2.* Absence of these two genes in *C. elegans* leads to a transverse spindle orientation in P1 [64]. However, in the *par*-*2/let*-*99* double mutant the majority of embryos orients the cleavage spindle longitudinally in both blastomeres (Rose and Kemphues, 1998). These genes are not only missing from the *D. coronatus* genome but are also absent in *Pristionchus and Panagrellus.* Since the latter show a *C. elegans-like* AB spindle orientation the molecular differences between *C. elegans* and *D. coronatus* can at most be considered a prerequisite for an alternative spindle orientation. The visible presence of a central cortical region in P1 and AB rather than in P1 alone has been suggested to be responsible for capturing spindle microtubules in both cells of *D. coronatus* resulting in a-p spindle orientation [11].

### Early transcriptome: species-specific orthologous clusters and their expression

To investigate to what extent the initial steps of embryogenesis in *D. coronatus* are reflected on the gene expression level, we sequenced 1-8-cell stages and compared their transcriptome with available data of similar stages from *C. elegans* [15] and *A. suum*[44]. By assembling transcriptomes from four independent biological replicates (Table S1), we retrieved in total more than 6,500 transcripts with a median length of 381 bp (Fig. S2, Table 3). For around 70% of these we could identify open reading frames (ORFs) allowing a successful inference of an early proteome.

We used our transcriptome data from *D. coronatus* to perform an orthology clustering between the three nematodes. We identified protein clusters that are shared among all three species as well as ones that are exclusively expressed in individual representatives during early development (Fig. 7). By subtracting orthologs expressed during early embryogenesis in *C. elegans*, we identified genes expressed only in the other two species. We retrieved 119 orthologous clusters shared between *D. coronatus* and *A. suum* as well as 350 (comprising nearly 1,500 genes) expressed exclusively in the early *D. coronatus* embryo (Fig. 7). While we found that orthologs of more than 60% of these genes are present in the genomes of *C. elegans* and *A. suum*, they are not transcribed during early embryogenesis in these species, suggesting interspecific heterochronic shifts of expression patterns. Exploring which potential functions these early expressed genes might have in *D. coronatus* more than 500 were classified as “unknown” as either no homology to any protein could be detected or homologous proteins are also unknown in their respective functions. In addition, we found over 1,300 *D. coronatus*-specific transcripts expressed as a single sequence only (“singletons”, Fig. 7).

**Fig. 7:**
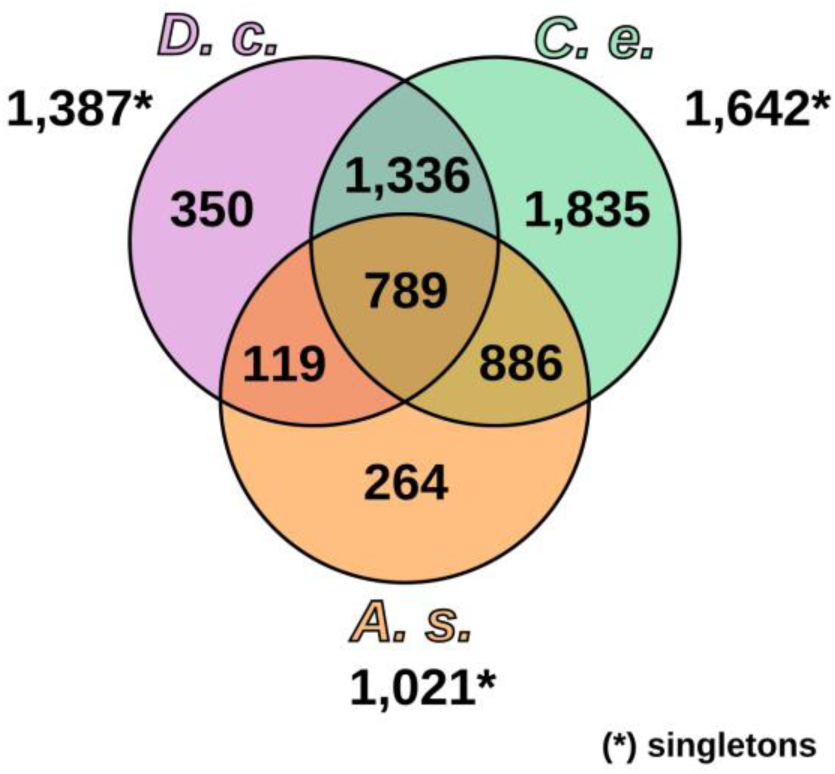
Early expressed clusters (protein families) of *D. coronatus* in comparison to the model *C. elegans* and the parasitic *A. suum.* For reference the number of singletons is given for each species. *C.e.*, *Caenorhabditis elegans*; *D.c.*, *Diploscapter coronatus*; *A.s.*, *Ascaris suum.*

We searched for significantly overrepresented gene ontology (GO) terms concerning genes specifically expressed in the early *D. coronatus* embryo (1-8 cell stage; Fig. 7) and in its whole genome. By far the most overrepresented was “regulation of centromere complex assembly” (GO:0090230; >200-fold). Related to this is “CENP-A containing nucleosome assembly at centromere” (GO:0034080; 48-fold). While it must be assumed that CENP-A is ubiquitously essential for mitosis, the lack of expression in *C. elegans* can be explained most easily with a maternal supply of the protein. Other potentially interesting overrepresented terms are “cytokinesis, initiation of separation” (GO:0007108; 64-fold) as well as chromatin remodeling associated terms such as “histone H2A acetylation” (GO:0043968; 48-fold) and “NuA4 histone acetyltransferase complex” (GO:0035267; 6-fold).

NuA4 is involved in the acetylation of H2A in yeast nucleosomes to exchange H2A for H2A.Z (Altaf et al., 2010) which in turn regulates gene expression. In the *C. elegans* embryo H2A.Z (or HTZ-1) is expressed in every blastomere and essential for normal development [65]. The observed NuA4-dependent acetylation in the early *D. coronatus* embryo is consistent with the assumption that massive zygotic transcription is required.

In search for further genes that could play a role for the unique *D. coronatus* early development, we looked for genes that are expressed in the early embryo but for which no orthologs were found in the genomes of *C. elegans* and *A. suum* and analyzed their expression. In this category, we detected less than 10 genes. As we could not retrieve orthologs of any of these genes in other reference systems like *Drosophila*, zebrafish or mouse, we are presently unable to speculate about their potential *Diploscapter-specific* functions.

As an alternative approach, we have started to look for protein domains significantly enriched in the early transcriptome of *D. coronatus* in comparison to *C. elegans* and *A. suum.* So far, we found a variety of enriched domains giving the chance to further investigate the role of certain proteins for developmental peculiarities in this species.

### The parthenogenetic *D. coronatus* shows high “allelic” divergence

Analysing the genome of *D. coronatus* we had observed an unexpected degree of heterozygosity in the high-quality draft genome (N50 = 1,007,652 bp, number of scaffolds = 511) resulting in the prediction of two “alleles” per gene in the Augustus pipeline. We observed a similar pattern when using Sanger sequencing methods on cloned PCR products of rRNA genes from individual worms. In fact, we retrieved several sequences per rRNA locus per individual (Figs. 8a and S1a, c). Aligning these sequences showed distinct single nucleotide polymorphisms (SNPs) and in a phylogenetic analysis all sequences could be allocated to one of two distinct” alleles” (Figs. 8b and S1b, d).

**Fig. 8:**
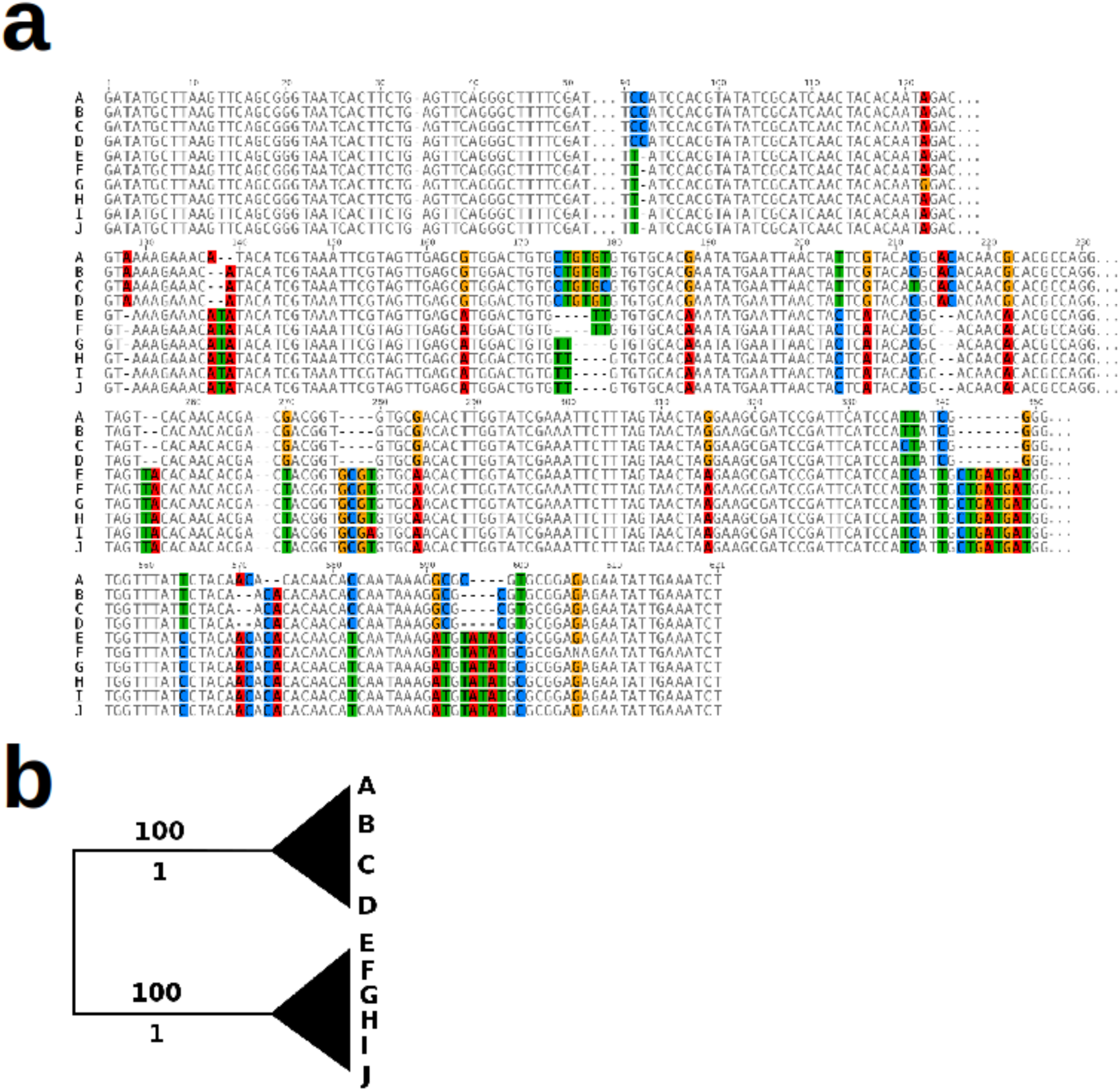
Sequence comparison of the small internal transcribed spacer (ITS) rDNA gene of *D. coronatus.* a, Sequence alignment of individual clones (A-J) shows selected regions with distinct single nucleotide polymorphisms (SNPs). b, Collapsed Maximum Likelihood tree representing clustering of sequenced clones. Bootstrap values are shown above and posterior probability beneath branches.

This was on a larger scale reflected in the orthology clustering we performed for this work. Here, we found more than 3,000 *D. coronatus*-specific clusters and more than 50% of these contained two in-paralogs (“doublets”), which is a multiple of what has been observed in the other studied nematodes (e.g. 2% in *C. elegans*; Fig. 1). In the complete genome 66% of all genes in *D. coronatus* were found to exist in doublets [2]. In contrast, the number of clusters consisting of a species-specific single protein (“singletons”) was by far the smallest in *D. coronatus* (2,727; *C. e.* 6,067).

*D. coronatus*-specific clusters comprising two proteins is in accordance with our finding that in this species most genes are present in two distinct “alleles” (see [2]). In contrast, in *C. elegans* we found only three such examples according to our definition (Table 2).

**Table 2.**
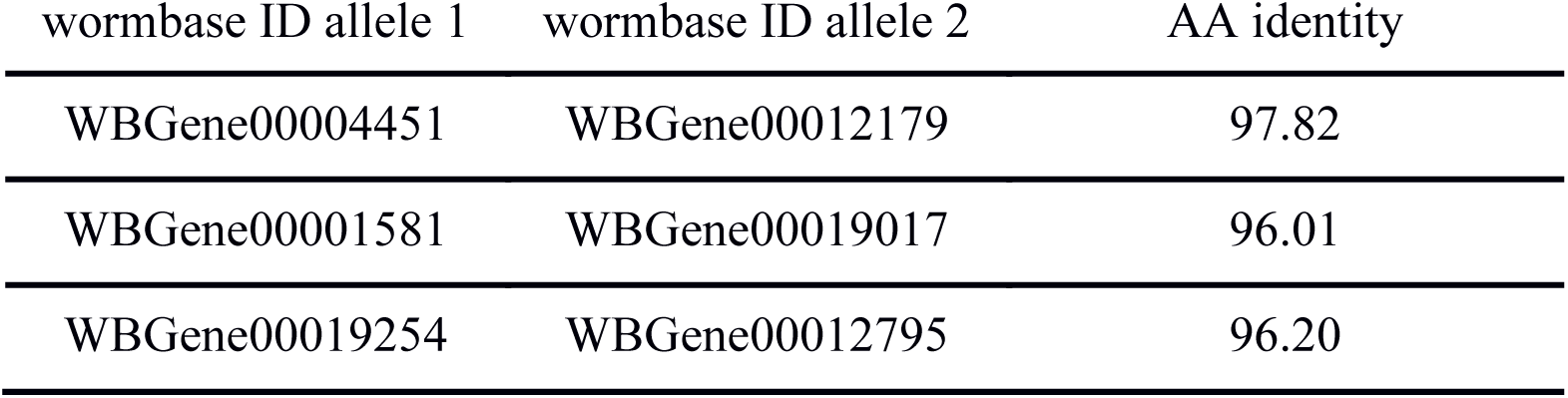
Genes existing in two distinct alleles identified in the *C. elegans* genome at a threshold of at least 95% AA identity.

| wormbase ID allele 1 | wormbase ID allele 2 | AA identity |
| --- | --- | --- |
| WBGene00004451 | WBGene00012179 | 97.82 |
| WBGene00001581 | WBGene00019017 | 96.01 |
| WBGene00019254 | WBGene00012795 | 96.20 |

**Table 3.**
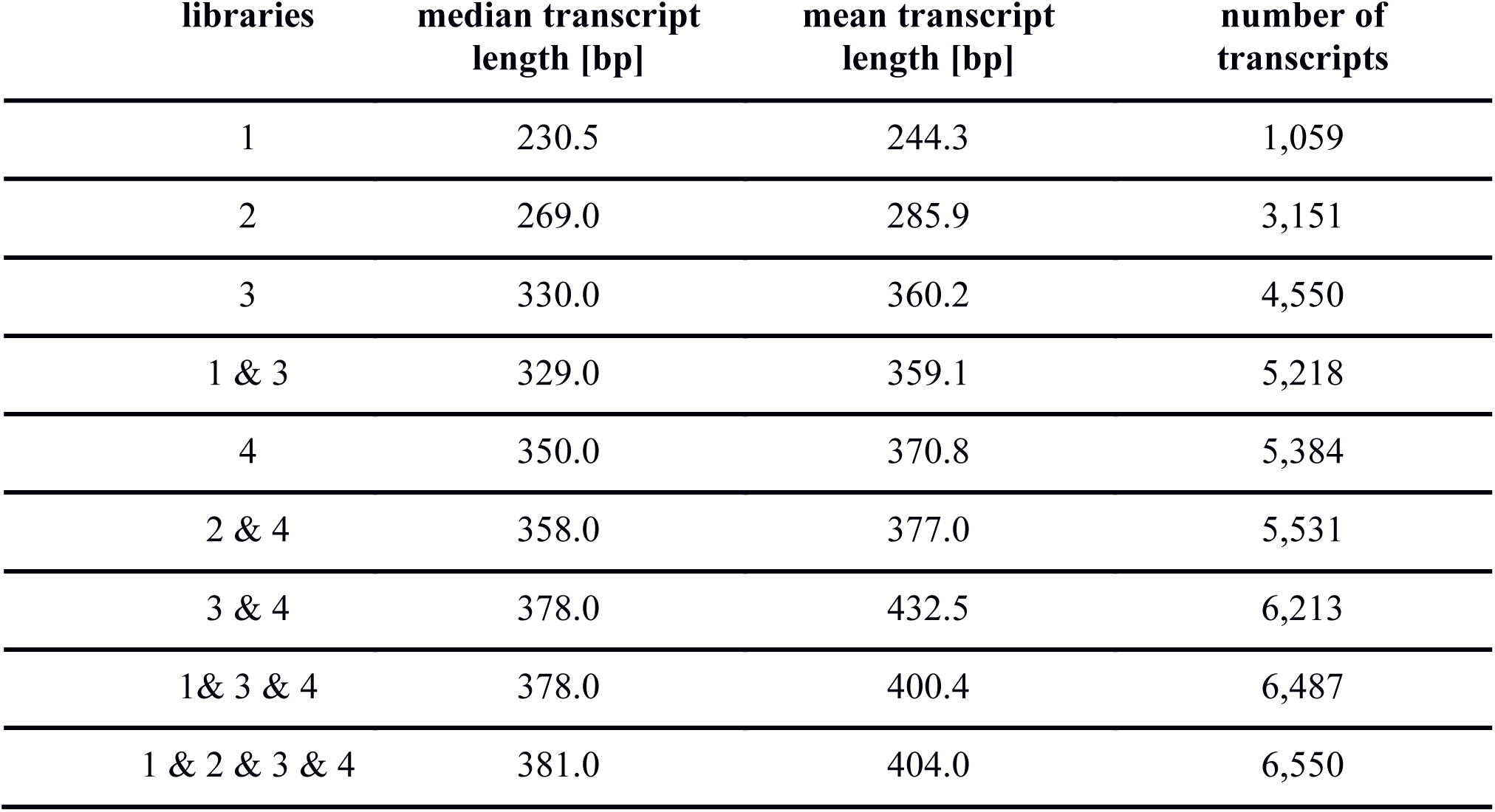
Average transcript lengths and numbers of transcripts of four independently sequenced
*D. coronatus* libraries for early developmental stages

| libraries | median transcript length [bp] | mean transcript length [bp] | number of transcripts |
| --- | --- | --- | --- |
| 1 | 230.5 | 244.3 | 1,059 |
| 2 | 269.0 | 285.9 | 3,151 |
| 3 | 330.0 | 360.2 | 4,550 |
| 1 & 3 | 329.0 | 359.1 | 5,218 |
| 4 | 350.0 | 370.8 | 5,384 |
| 2 & 4 | 358.0 | 377.0 | 5,531 |
| 3 & 4 | 378.0 | 432.5 | 6,213 |
| 1 & 3 & 4 | 378.0 | 400.4 | 6,487 |
| 1 & 2 & 3 & 4 | 381.0 | 404.0 | 6,550 |

This pattern can be explained in two ways: (i) the independent accumulation of SNPs in non-recombining alleles (known as Meselson effect) in an old parthenogenetic lineage [66,67] or (ii) a hybrid origin of the parthenogenetic strain, where distinct alleles are inherited from the ancestral sexual species and genomic heterozygosity is maintained.

To investigate to what extent an accumulation of mutations occurred in the *D. coronatus* genome, we compared conserved single copy genes in the nematode phylum. We found that in *D. coronatus* each of these single copy genes exists in two distinct “alleles”. The number of AA differences between the two *D. coronatus* “alleles” is similar to AA differences between *C. elegans* and *C. remanei* (Fig. 9a). Calculation of dN/dS ratios in 11 arbitrarily selected single copy genes revealed a median value of 0.158 (Table S2) indicating negative selection. To identify potential differences between conserved and non-conserved protein domains we applied InterProScan [49]. As expected we found the number of non-synonymous exchanges to be lower than synonymous exchanges. But the ratio of synonymous vs. non-synonymous substitutions was again not significantly different when comparing the two *D. coronatus* “alleles” with the respective *C. elegans* vs. *C. remanei* orthologs (Fig. 9b). This is in line with (ii; see above), and can be easiest explained with a recent interspecies hybridization event.

**Fig. 9:**
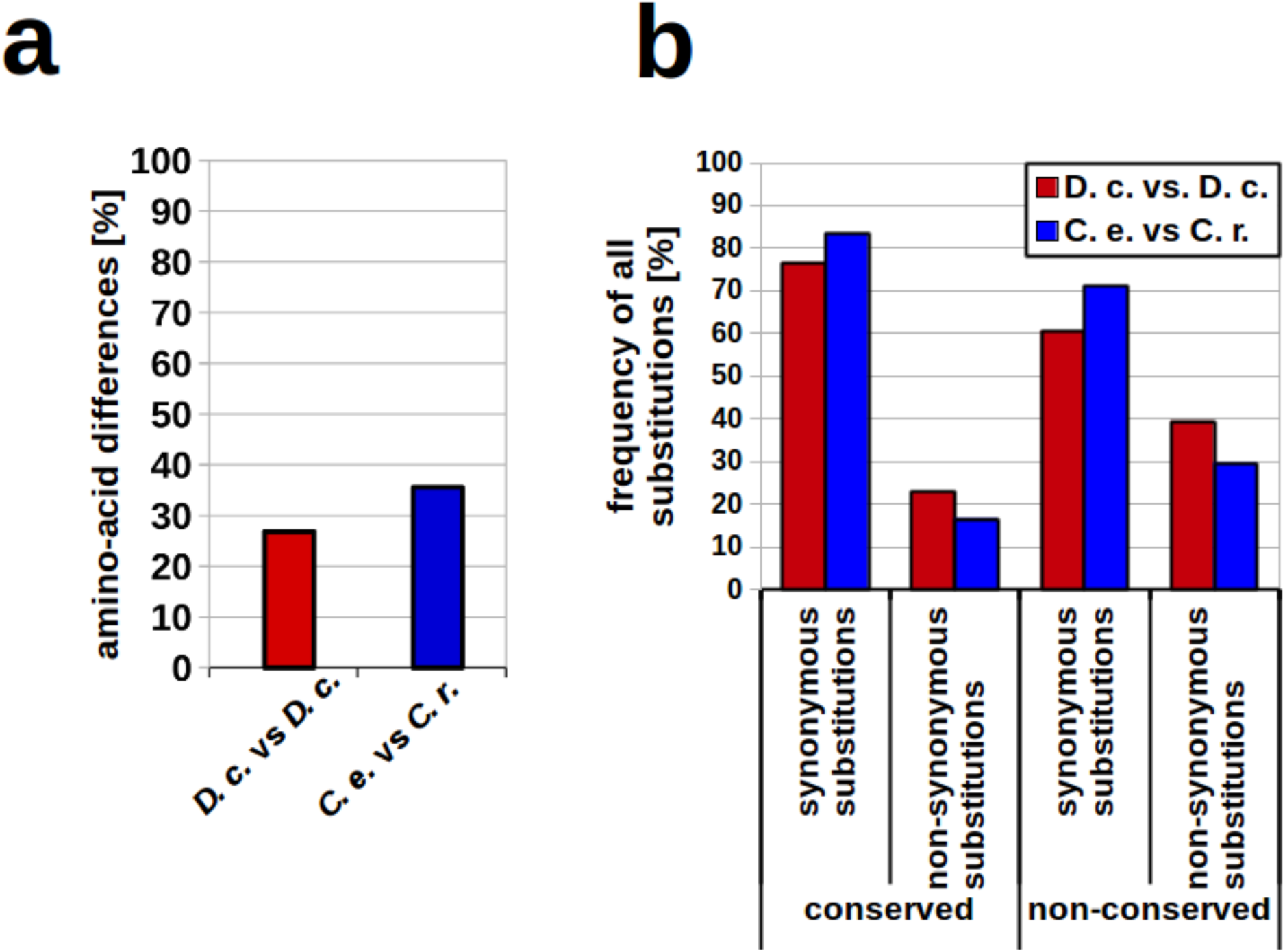
Analysis of *Diploscapter coronatus* genes. a, AA differences among alleles of single copy genes of *C. elegans* vs. *C. remanei* (blue; n=229) and the two “alleles” of *D. coronatus* orthologs (red; n=307); b, Percentage of synonymous and non-synonymous substitutions in non-conserved and conserved protein regions. Abbreviations: *C.e*, *C. elegans*; *C.r.*, *C. remanei*; *D.c*, *D. coronatus.*

### Differential expression of alleles

Incipient hybrids may face dosage compensation issues and proteins built from alleles inherited from different genomes might be incompatible or less efficient in molecular machineries. It is therefore possible that hybrid species need to silence one of the parental genomes [68]. Making use of the high-quality *D. coronatus* genome and our RNA-Seq data covering early embryogenesis we asked whether transcripts of one or both “alleles” are generated by screening for single nucleotide polymorphisms (SNPs) in comparison to an EST library. We found that in the vast majority of genes both “alleles” are expressed (Fig. S2).

## Discussion

Our previous studies of the parthenogenetic nematode *Diploscapter coronatus* focused on early embryogenesis [1,11] and on molecular regulators important for the oocyte-to-embryo transition [14]. Recently, we sequenced and assembled the genome of *D. coronatus* and here use these data as a reference to re-visit these questions in a more comprehensive scope.

### Meiosis, *D. coronatus-specific* genes and preservation of heterozygosity

Previously, it has been shown that *D. coronatus* passes through a truncated meiosis generating only one polar body [1]. We find that the gonad of *D. coronatus* differs in several aspects from *C. elegans* (Fig. 3a, b). The uniformity of germ cells suggests that due to the small size of the gonad the distal tip cell (dtc) prevents entrance into meiosis all along the gonadal tube (in contrast to *C. elegans*, see [69]), such that only the most proximal oocyte, possibly due to its translocation into the uterus, can escape its influence. Another conclusion is that there is no meiotic arrest of germ cells in the gonad as found in *C. elegans* [53], Kim:2013vx} and therefore no sperm is needed to lift it.

Compared to *C. elegans*, *D. coronatus* seems to follow a different strategy for the control of germ cell development. It would be attractive to study in this respect the role of the dtc in nematodes with particularly small gonads and a low brood size like *D. coronatus* or the parthenogenetic *Plectus sambesii* [70] and match it with representatives possessing extremely long gonadal arms, like *Ascaris*, producing millions of eggs [71].

The fact that no orthologs were found of crucial genes required for the generation of the synaptonemal complex (e. g. *syp*-*1/*-*2/*-*3*) and chromosome-specific adapters which are also involved in proper meiosis (*zim-1/*-*2/*-*3*, *him*-*8*; also see [2]) could mean that no crossing-over and thus no recombination takes place. This is consistent with our inability to detect paired meiotic chromosomes (see also [72]). However, the finding that the genes listed above are absent in *Pristionchus* and *Panagrellus* as well (Fig. 2), does not offer a straight forward explanation. The genetic control of meiosis (and other processes) seems to differ generally between *Caenorhabditis* and other nematodes and may involve other genes across the phylum.

We identified more than 3,300 *D. coronatus*-specific clusters comprising more than 7,500 genes in its genome. However, the role of most of these remains elusive since no orthologs have been found in other model systems (see Results and Fig. 1). Looking at early expressed genes alone we recovered more than 500 genes of unknown function (see Results and Fig. 7).

A possible explanation for how the heterozygosity in *D. coronatus* can be preserved while passing through just one meiotic division, including the separation of chromatids rather than homologous chromosomes during meiosis I (“inverted meiosis”; [73]) has been discussed in [2].

### Polarity, asymmetry and absence of orthologs

Microscopical analysis of early embryogensis in *D. coronatus* revealed certain idiosyncrasies [1,11]. Here we show that the process of initial polarity establishment during the 1-cell stage differs markedly from *C. elegans* (Figs. 4 and 5). Based on our ortholog clustering for the *D. coronatus* genome and eight other nematode genomes (Fig. 1) we conclude that certain genes crucial for early embryogenesis in *C. elegans* are absent in *D. coronatus*, *P. pacificus*, *P. redivivus* and *A. suum* (Fig. 2). With respect to polarity establishment, we did not find orthologs of essential genes known from *C. elegans*, such as *gpr*-*1/*-*2* in these species indicating differences in the establishment of polarity. This appears particularly plausible for the parthenogenetic *D. coronatus*, where sperm as initial trigger is missing and where orientation of the anterior-posterior egg axis is obviously specified in a random fashion [1]. However, by scanning the *D. coronatus* genome for known GoLoco domain proteins involved in mitosis in vertebrates we found that the human GPR-1/2 homologue LGN [57] has orthologs in *D. coronatus* (Fig. S3). This suggests that *C. elegans* acquired new adaptor proteins for division while *D. coronatus* relies on the established set of proteins known from outgroup species. It remains to be determined whether LGN functionally replaces GPR-1/2 in the *D. coronatus* 1-cell stage and to what extent the modified dcLET-99 homolog (see Results and Fig. 4) is involved in the establishment of early polarity.

Looking at conserved protein complexes which are essential for maintaining already established polarity, such as PAR-3/PAR-6/PKC-3, or PAR-1/-2 [60], we found respective orthologs in *D. coronatus*, except for PAR-2 (Fig. 2). This result is in accordance with earlier studies, where it was proposed that the PAR-2/-3 system known from *C. elegans* evolved specifically in the genus *Caenorhabditis* [8]. What could be an alternative mechanism for establishing polarity in other nematodes like *D. coronatus*? It was shown earlier that in *C. elegans* the *par*-*2* function can be replaced by *lgl*-*1* (ortholog of the tumor surpressor gene *lethal giant larvae*) if it is overexpressed [74]. *lgl* is known from various animal systems as a regulator of asymmetric cell division [75] indicating its high conservation. Its presence and the simultaneous absence of *par*-*2* in *D. coronatus* suggest an ancestral molecular mechanism of asymmetric cleavage there and in the other non-*Caenorhabditis* species studied, as known from animals outside the nematodes [76,77].

Another idiosyncracy in *D. coronatus* development is that the spindle in AB performs the same rotation as in P_1_ resulting in its a-p orientation. In *C. elegans* spindle microtubules in P_1_ seem to compete for attachment to a bleb-like site at the anterior cortex [78,79]. A prominent clear cortical region in the cortex of both 2-cell blastomeres in *D. coronatus* may indicate a symmetric distribution of components responsible for capturing spindle microtubules[11]. In the more basal nematode *Romanomermis culicivorax* (clade 2) a comparable effect on spindle behavior appears to be exerted in the 2-cell stage by the “region of the first midbody”. Its ablation results in a perpendicular spindle orientation [80]. More data on the gene regulatory networks and intracellular constraints in *D. coronatus* should help us to find the molecular basis for the aberrant orientation of the cleavage spindle in AB. Presently, we speculate that here an original mechanism has been replaced in more derived nematodes, including *C. elegans*, while the phylogenetic branch comprising *Diploscapter* and *Protorhabditis* [4,10,81], constitutes an atavism.

### GO term analysis

We performed a gene ontology analysis for the “early transcriptome” and found over 70 terms overrepresented specifically in *D. coronatus* (data not shown). It appears likely that enriched terms inform about important underlying biological processes [82]. While this approach should increase the likelihood for identifying such essential events it can only play an advisory role in finding the most relevant, enriched annotation terms [83].

Many of the highly overrepresented terms are associated with chromosomal function behavior in *D. coronatus*, for instance concerning centrosomes, chromosome structure, telomeres, DNA replication and gene regulation. This is consistent with the view that in *D. coronatus* these aspects differ considerably from *C. elegans*.

### Origin of parthenogenesis and reduction of chromosomes

Different mechanisms and conditions have been described that could lead to the establishment of parthenogenesis [3,84,85].

In the parthenogenetic *D. coronatus* where neither males nor sperm have been observed but a high degree of heterozygosity persists, we conceive two possible scenarios. One is hybridization of closely related gonochoristic species followed by the evolution of parthenogenesis and the other a spontaneous origin of parthenogenetic reproduction followed by independent accumulation of mutations in alleles. In the latter case the dN/dS ratio in the studied single copy genes should be high as both non-synonymous and synonymous mutations are expected to accumulate to a similar degree in either of the two alleles. In contrast, a recent interspecies hybridization event should still show the footprints of purifying (negative) selection acting in the sexually reproducing parent species [86].

Looking at the AA sequences of arbitrarily selected single copy genes we found a high level of heterozygosity similar to other parthenogenetic species resulting from crosses between two bisexual partners. The percentage of AA exchanges in these *D. coronatus* genes were not significantly different from those found in two *Caenorhabditis* species (Fig. 9) indicating similar conservation of both genomic variants.

The high genomic heterozygosity with two distinguishable “alleles”s per gene, including the here described rDNA genes (Figs. 8, S1) and single-copy genes (Fig. 9), plus the low dN/dS ratio described above, can be easiest explained with a recent event of interspecies hybridization between two closely related representatives where each parental genome has still preserved a major part of its ancestral functionality. In fact, this has recently been suggested for more distantly related parasitic nematodes of the genus *Meloidogyne* [68,87,88]) and also suggested for parthenogens of the phylogenetically closer genus *Panagrolaimus* (bioRxiv: [89]).

*D. coronatus* possesses only a single chromosome in the haploid set while *C. elegans* and most other studied free-living nematode species of clades 9-12 contain 6 chromosomes or more [90,91] (our unpublished data). However, a close relative of *Diploscapter*, *Protorhabditis* sp. (laboratory strains JB 122), also contains just a single chromosome while other members of this genus have six like *C. elegans* (E.S., unpublished data). The most parsimonious explanation for this minimal number in selected species is a comprehensive chromosome fusion. While a reduction in chromosome number due to fusion has been described in a variety of cases [92,93], integration of all chromosomes into a single one would be a very extraordinary case deserving further attention. A combined detailed phylogenetic and chromosome analysis may reveal whether the assumed fusion has occurred once in a common ancestor or several times independently, whether this has been a stepwise modification or a single total fusion event, and how original chromosomes are arranged in this new construct. While we can only speculate about a possible mechanism for such a dramatic event, it may be worthwhile to investigate whether each chromosome in *D. coronatus* represents a complete haploid parental genome which has remained intact and functional due to the absence of recombination. In this case chromosome fusion would have taken place most likely prior to the envisaged interspecies hybridization.

A haploid chromosome number of n=1 is neither a necessary prerequisite for parthenogenesis in nematodes nor a consequence of it, since in the bisexual *Ascaris* (n=2) a variant exists with n=1 (*var. univalens*; {Beneden:7uHQzYCc}) and in the parthenogenetic species *Acrobeloides nanus* and *Plectus sambesii* we counted n=6.

It is an exciting question why in contrast to the closely related genera *Diploscapter* and *Protorhabditis* no parthenogenetic representatives have been found among the around 50 *Caenorhabditis* species isolated so far [94] (NCBI Taxonomy). Taking into account the many idiosyncrasies of the taxon *Caenorhabditis* concerning the control of development (see e.g. Fig. 2) it seems not unlikely that its molecular circuitry, allowing for instance a particularly rapid propagation, has not allowed the establishment of parthenogenesis.

### The genus *Caenorhabditis* vs. *Diploscapter* and other nematodes

Many *C. elegans* genes not found in *D. coronatus* were also absent from other non-*Caenorhabditis* species (Fig. 2). The peculiarities in early development of *D. coronatus* can thus not be explained just by the absence of these genes. It is feasible that in *Diploscapter* certain processes differ from *C. elegans* due to its parthenogenetic mode of reproduction. However, in light of the genomic similarities all studied non-*Caenorhabditis* nematodes may use the same alternative pathways (or at least alternative components) to control essentially identical developmental processes during oogenesis and early embryogenesis (Developmental System Drift [7]). Therefore, it appears more likely that these differences gave the freedom to establish parthenogenesis while preventing it in the genus *Caenorhabditis.* So far only part of the identified differences on the molecular level can be correlated to the described variations on the cellular level where “many roads lead to Rome” [95,96]. Future studies have to reveal to what extent the developmental characteristics of *D. coronatus* can be explained with variations on the transcriptional and post-transcriptional level.

Even if with improved methods in the one or other case a credible ortholog of a *C. elegans* gene could be excavated that we here counted as being absent with our approach in the non-*Caenorhabditis* representatives, the differences on the level of genes and gene products in comparison to *Diploscapter* (and the other studied nematodes) will still remain remarkable. The particularly rapid evolutionary diversification in the genus *Caenorhabditis* [4] may be related to multifaceted opportunistic life styles. In permanently changing environments this allows on the one hand short generation times and large progenies whenever food is available in abundance and on the other hand long-term survival under harsh conditions. Extended studies on additional *Caenorhabditis* species and close outgroups will have to reveal whether this genus is really as uniform with respect to its control of development as our limited studies suggest and at which branching points in the phylogenetic tree novelties arose. If the methodology to establish a transcriptional lineage of the early *C. elegans* embryo [97] is applicable to *D. coronatus* and other nematodes as well, evolutionary changes in time and space could even be traced on a single cell level.

## Conclusions

The parthenogenetic *D. coronatus* reveals a variety of differences compared to *C. elegans* on the level of cells, chromosomes, genome and transcriptome indicating alternative routes for nematode development and reproduction. Thus, it appears to be an attractive study object to better understand the intricate pathways of evolutionary change among closely related species. Our comparative study further supports the view that the genus *Caenorhabditis* cannot be taken as a blueprint for the genetic control of developmental and reproductive processes in nematodes as it shows a number of idiosyncrasies absent in the other studied representatives. Future avenues to follow in order to reveal further developmentally relevant differences between *D. coronatus*, *C. elegans* and other rhabditid nematodes could be: (i) the role of early transcription vs. maternal supply, (ii) structure and function of the single chromosome (n=1) in *D. coronatus* (Fig. 3c) assumed to be the result of fusion, (iii) meiotic pairing and crossover, apparently absent in *D. coronatus*, (iv) the mechanism of chromosome separation.

## Legends for Suppplementary Figures

**Fig. S1:**
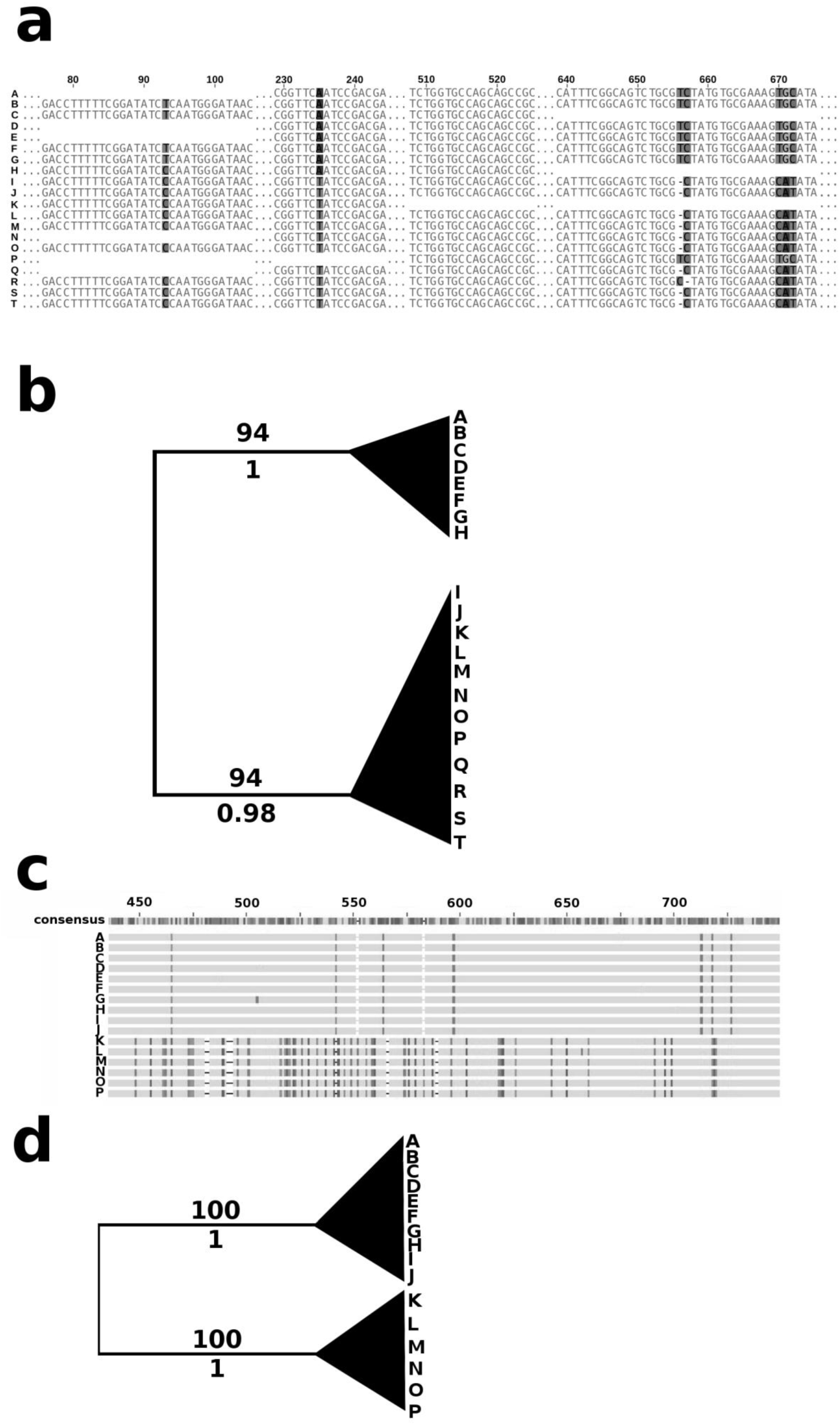
a, c, Sequence comparison of the small (SSU) and large subunit (LSU) rDNA genes of *D. coronatus.* b, Collapsed Maximum Likelihood (ML) tree representing clustering of sequenced clones for the SSU rDNA gene. (d) Collapsed ML tree representing clustering of sequenced clones for the LSU rDNA gene. Bootstrap values are shown above and posterior probability values beneath branches.

**Fig. S2:**
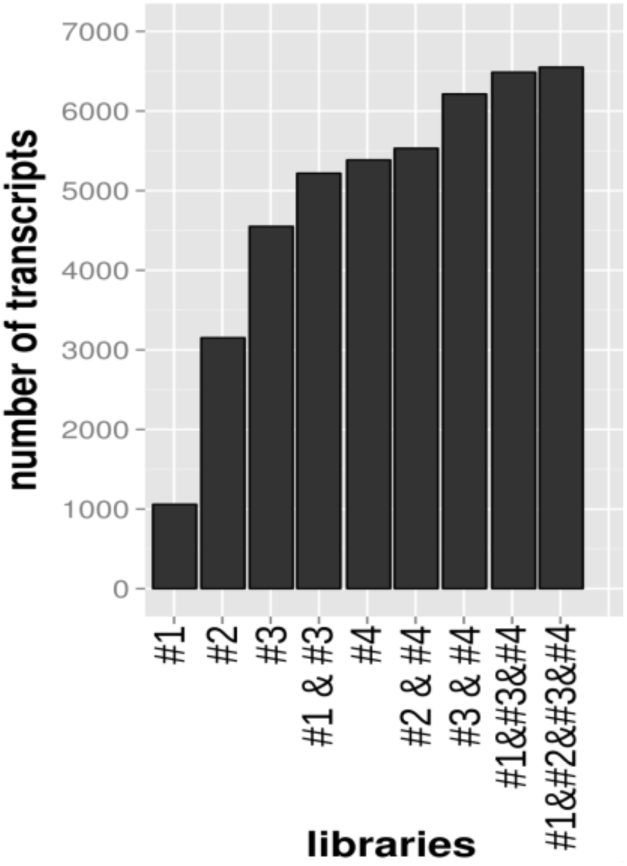
Binning of different combinations of replicates. Combining all four replicates the numbers of expressed sequences appear to saturate at about 6,500 transcripts (Table 3).

**Fig. S3:**
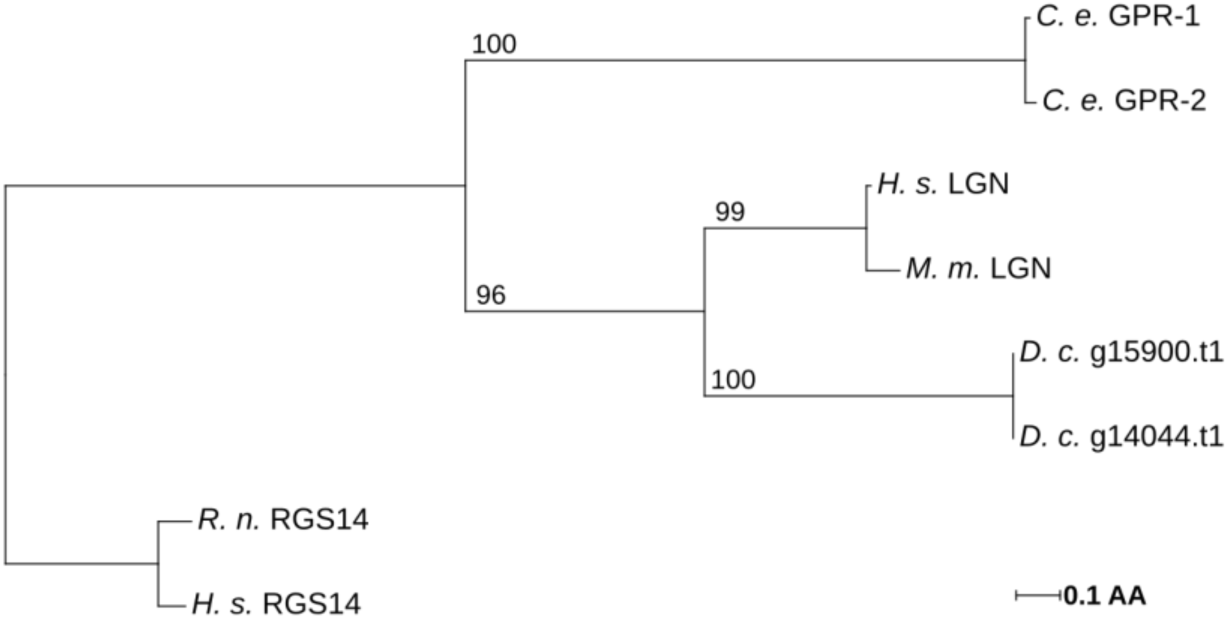
Phylogenetic tree representing GoLoco (Pfam ID PF02188) domain proteins of *D. coronatus* (*D. c.*), *C. elegans* (*C. e.*), human (*H. s.*), rat (*R. n.*) and mouse (*M. m.*).

## Supplementary tables

**Table S1:** Retrieved paired-end reads from RNA sequencing of four independent *D. coronatus* samples

| Library<sup>a</sup> | paired-end reads | read length [bp] | discarded paired-end reads<sup>b</sup> | successfully mapped reads<sup>c</sup> | assembled and mapped transcripts | Illumina platform |
| --- | --- | --- | --- | --- | --- | --- |
| #1 | 6,979,616 | 75 | 3,402,448 | 173,936 (2.5%) | 1,059 | MiSeq |
| #2 | 8,898,677 | 75 | 7,947,398 | 673,291 (7.6%) | 3,151 | MiSeq |
| #3 | 31,199,716 | 100 | 1,111,574 | 1,779,470 (6.0%) | 4,550 | HiSeq |
| #4 | 28,416,086 | 100 | 1,341,994 | 1,349,244 (5.0%) | 5,384 | HiSeq |

a, each sample >100 early embryos (1- to 8-cell stages); b, paired-end reads with low sequencing quality for the 5'- and/or 3'-ends were removed; c, successfully mapped raw reads to the *D. coronatus* EST library and transcriptome.

**Table S2:** Ratio of non-synonymous to synonymous base exchanges in 11 conserved single copy genes of *D. coronatus.*

| Fosmid ID* | *C.e.* gene name | dN/dS |
| --- | --- | --- |
| C09G12.9 | *tsg-101* | 0.096 |
| F08D12.1 | *srpa-72* | 0.144 |
| F21H11.2 | *sax-2* | 0.842 |
| F32E10.1 | *nol-10* | 0.129 |
| F39C12.1 | F39C12.1 | 0.388 |
| F41E6.9 | *vps-60* | 0.022 |
| F44A6.1 | *nucb-1* | 0.093 |
| F48F5.5 | *fce-2* | 0.337 |
| F53G2.6 | *tsr-1* | 0.158 |
| Y47D3A.28 | *mcm-10* | 0.284 |
| Y65B4A.1 | Y65B4A.1 | 0.277 |
| median |  | 0.158 |

\*orthologs taken from Mitreva et al., 2011

## References

1. Lahl V, Sadler B, Schierenberg E. Egg development in parthenogenetic nematodes: variations in meiosis and axis formation. Int J Dev Biol. 2006;50:393–7.

2. Hiraki H, Kagoshima H, Kraus C, Schiffer PH, Ueta Y, Kroiher M, et al. Genome analysis of Diploscapter coronatus: insights into molecular peculiarities of a nematode with parthenogenetic reproduction. BMC Genomics. 2017;18:478.

3. Simon J, Delmotte F, Rispe C, Crease T. Phylogenetic relationships between parthenogens and their sexual relatives: the possible routes to parthenogenesis in animals. Biol J Linn Soc. 2003;79:151–63.

4. Kiontke K, Gavin N, Raynes Y, Roehrig C, Piano F, Fitch D. Caenorhabditis phylogeny predicts convergence of hermaphroditism and extensive intron loss. Proceedings of the National Academy of Sciences. 2004;101:9003.

5. Kiontke KC, Félix M-A, Ailion M, Rockman MV, Braendle C, Penigault J-B, et al. A phylogeny and molecular barcodes for Caenorhabditis, with numerous new species from rotting fruits. BMC Evol Biol. 2011;11:339.

6. Schierenberg E, Sommer R. Reproduction and development in nematodes. Schmidt-Rhaesa A, editor. Handbook of Zoology. Berlin: De Gruyter; 2014.

7. True JR, Haag ES. Developmental system drift and flexibility in evolutionary trajectories. Evol Dev. 2001;3:109–19.

8. Schiffer PH, Kroiher M, Kraus C, Koutsovoulos GD, Kumar S, R Camps JI, et al. The genome of Romanomermis culicivorax: revealing fundamental changes in the core developmental genetic toolkit in Nematoda. BMC Genomics. BioMed Central; 2013;14:923.

9. Schiffer PH, Nsah NA, Grotehusmann H, Kroiher M, Loer C, Schierenberg E. Developmental variations among Panagrolaimid nematodes indicate developmental system drift within a small taxonomic unit. Development Genes and Evolution. 2014;224:183–8.

10. Kiontke K, Fitch D. The phylogenetic relationships of Caenorhabditis and other rhabditids. 2005.

11. Lahl V, Schulze J, Schierenberg E. Differences in embryonic pattern formation between Caenorhabditis elegans and its close parthenogenetic relative Diploscapter coronatus. Int J Dev Biol. 2009;53:507–15.

12. Miller MA, Ruest PJ, Kosinski M, Hanks SK, Greenstein D. An Eph receptor sperm-sensing control mechanism for oocyte meiotic maturation in Caenorhabditis elegans. Gene Dev. Cold Spring Harbor Lab; 2003;17:187–200.

13. Miller M, Nguyen V, Lee M, Kosinski M, Schedl T, Caprioli R, et al. A sperm cytoskeletal protein that signals oocyte meiotic maturation and ovulation. 2001;291:2144.

14. Heger P, Kroiher M, Ndifon N, Schierenberg E. Conservation of MAP kinase activity and MSP genes in parthenogenetic nematodes. Dev Biol. 2010; 10:51.

15. Hashimshony T, Wagner F, Sher N, Yanai I. CEL-Seq: Single-Cell RNA-Seq by Multiplexed Linear Amplification. CellReports. The Authors; 2012;2:1–8.

16. Wang J, Mitreva M, Berriman M, Thorne A, Magrini V, Koutsovoulos G, et al. Silencing of Germline-Expressed Genes by DNA Elimination in Somatic Cells. Developmental Cell. Elsevier Inc; 2012;23:1–9.

17. Neaves WB, Baumann P. Unisexual reproduction among vertebrates. Trends in Genetics. 2011;27:81–8.

18. Brenner S. The genetics of Caenorhabditis elegans. Genetics. Genetics Society of America; 1974;77:71–94.

19. Schnabel R, Hutter H, Moerman D, Schnabel H. Assessing Normal Embryogenesis inCaenorhabditis elegansUsing a 4D Microscope: Variability of Development and Regional Specification. Dev Biol. 1997;184:234–65.

20. Williams BD, Schrank B, Huynh C, Shownkeen R, Waterston RH. A genetic mapping system in Caenorhabditis elegans based on polymorphic sequence-tagged sites. Genetics. Genetics; 1992;131:609–24.

21. Holterman M, Holovachov O, van den Elsen S, van Megen H, Bongers T, Bakker J, et al. Small subunit ribosomal DNA-based phylogeny of basal Chromadoria (Nematoda) suggests that transitions from marine to terrestrial habitats (and vice versa) require relatively simple adaptations. Mol Phylogenet Evol. 2008;48:758–63.

22. Floyd R, Abebe E, Papert A, Blaxter M. Molecular barcodes for soil nematode identification. Mol Ecol. 2002;11:839–50.

23. Sonnenberg R, Nolte A, Tautz D. An evaluation of LSU rDNA D 1-D 2 sequences for their use in species identification. Frontiers in Zoology. 2007;4:6.

24. Vrain TC. Restriction Fragment Length Polymorphism Separates Species of the Xiphinema americanum Group. J Nematol. Society of Nematologists; 1993;25:361–4.

25. Stamatakis A. RAxML-VI-HPC: maximum likelihood-based phylogenetic analyses with thousands of taxa and mixed models. Bioinformatics. 2006;22:2688–90.

26. Li L, Stoeckert CJ, Roos DS. OrthoMCL: identification of ortholog groups for eukaryotic genomes. Genome Res. 2003;13:2178–89.

27. C. elegans Sequencing Consortium. Genome sequence of the nematode C. elegans: a platform for investigating biology. Science. 1998;282:2012–8.

28. Stein LD, Bao Z, Blasiar D, Blumenthal T, Brent MR, Chen N, et al. The genome sequence of Caenorhabditis briggsae: a platform for comparative genomics. PLoS Biol. 2003;1:E45.

29. Mortazavi A, Schwarz EM, Williams B, Schaeffer L, Antoshechkin I, Wold BJ, et al. Scaffolding a Caenorhabditis nematode genome with RNA-seq. Genome Res. 2010;20:1740–7.

30. Dieterich C, Clifton SW, Schuster LN, Chinwalla A, Delehaunty K, Dinkelacker I, et al. The Pristionchus pacificus genome provides a unique perspective on nematode lifestyle and parasitism. Nature Genetics. 2008;40:1193–8.

31. Srinivasan J, Dillman AR, Macchietto MG, Heikkinen L, Lakso M, Fracchia KM, et al. The draft genome and transcriptome of Panagrellus redivivus are shaped by the harsh demands of a free-living lifestyle. Genetics. Genetics Society of America; 2013;193:1279–95.

32. Jex AR, Liu S, Li B, Young ND, Hall RS, Li Y, et al. Ascaris suum draft genome. Nature. Nature Publishing Group; 2011;479:1–8.

33. Blüthgen N, Brand K, Cajavec B, Swat M, Herzel H, Beule D. Biological profiling of gene groups utilizing Gene Ontology. Genome Inform. 2005;16:106–15.

34. Conesa A, Götz S, García-Gómez JM, Terol J, Talón M, Robles M. Blast2GO: a universal tool for annotation, visualization and analysis in functional genomics research. Bioinformatics. 2005;21:3674–6.

35. Holterman M, van der Wurff A, van den Elsen S, van Megen H, Bongers T, Holovachov O, et al. Phylum-wide analysis of SSU rDNA reveals deep phylogenetic relationships among nematodes and accelerated evolution toward crown clades. Mol Biol Evol. 2006;23:1792–800.

36. De Ley P, Blaxter ML. Systematic position and phylogeny. The Biology of Nematodes. The biology of nematodes; 2002. pp. 1–30.

37. Sievers F, Wilm A, Dineen D, Gibson TJ, Karplus K, Li W, et al. Fast, scalable generation of high-quality protein multiple sequence alignments using Clustal Omega. Mol Syst Biol. Nature Publishing Group; 2011;7:1–6.

38. Finn RD, Coggill P, Eberhardt RY, Eddy SR, Mistry J, Mitchell AL, et al. The Pfam protein families database: towards a more sustainable future. Nucleic Acids Res. 2016;44:D279–85.

39. Darriba D, Taboada GL, Doallo R, Posada D. ProtTest 3: fast selection of best-fit models of protein evolution. Bioinformatics. 2011;27:1164–5.

40. Baugh LR, Hill AA, Brown EL, Hunter CP. Quantitative analysis of mRNA amplification by in vitro transcription. Nucleic Acids Res. Oxford University Press; 2001;29:E29.

41. Bolger AM, Lohse M, Usadel B. Trimmomatic: a flexible trimmer for Illumina sequence data. Bioinformatics. Oxford University Press; 2014;30:2114–20.

42. Grabherr MG, Haas BJ, Yassour M, Levin JZ, Thompson DA, Amit I, et al. Full-length transcriptome assembly from RNA-Seq data without a reference genome. Nat Biotechnol. 2011;29:644–52.

43. Ben Langmead, Salzberg SL. Fast gapped-read alignment with Bowtie 2. Nat Meth. Nature Research; 2012;9:357–9.

44. Wang J, Garrey J, Davis RE. Transcription in pronuclei and one- to four-cell embryos drives early development in a nematode. Curr. Biol. 2014;24:124–33.

45. Li W, Godzik A. Cd-hit: a fast program for clustering and comparing large sets of protein or nucleotide sequences. Bioinformatics. 2006;22:1658–9.

46. Altschul SF, Gish W, Miller W, Myers EW. Basic local alignment search tool. Journal of Molecular Biology. 1990;215:403–10.

47. Mitreva M, Jasmer DP, Zarlenga DS, Wang Z, Abubucker S, Martin J, et al. The draft genome of the parasitic nematode Trichinella spiralis. Nature Genetics. Nature Publishing Group; 2011;43:228–35.

48. Thompson JD, Higgins DG, Gibson TJ. CLUSTAL W: improving the sensitivity of progressive multiple sequence alignment through sequence weighting, position-specific gap penalties and weight matrix choice. Nucleic Acids Res. Oxford University Press; 1994;22:4673–80.

49. Quevillon E, Silventoinen V, Pillai S, Harte N, Mulder N, Apweiler R, et al. InterProScan: protein domains identifier. Nucleic Acids Res. 2005;33:W116–20.

50. Zhang Z. KaKs_Calculator Manual. 2006;:1–10.

51. Li H, Handsaker B, Wysoker A, Fennell T, Ruan J, Homer N, et al. The Sequence Alignment/Map format and SAMtools. Bioinformatics. 2009;25:2078–9.

52. Danecek P, Auton A, Abecasis G, Albers CA, Banks E, DePristo MA, et al. The variant call format and VCFtools. Bioinformatics. Oxford University Press; 2011;27:2156–8.

53. Greenstein D. Control of oocyte meiotic maturation and fertilization. 2005.

54. Kimble J, Crittenden S. Germline proliferation and its control (August 15, 2005), WormBook, ed. The C. elegans Research Community, WormBook, doi/10.1895/wormbook. 1.13. 1. 2005.

55. Hechler HC. Postembryonic development and reproduction in Diploscapter coronatus (Nematoda, Rhabditidae). Helminth. Soc. Washington. 1968. pp. 24–30.

56. Albertson D, Rose AM, Villeneuve AM. Chromosome organization, mitosis, meiosis. The nematode C. elegans, II. Cold Spring Harbor Laboratory Press; 1997.

57. Gonczy P, Rose LS. Asymmetric cell division and axis formation in the embryo. 2005;:1–20.

58. Cowan CR, Hyman AA. Centrosomes direct cell polarity independently of microtubule assembly in C. elegans embryos. Nature. 2004.

59. Tsou M-FB, Hayashi A, DeBella LR, McGrath G, Rose LS. LET-99 determines spindle position and is asymmetrically enriched in response to PAR polarity cues in <i>C. elegans</i> embryos. Development. The Company of Biologists Ltd; 2002;129:4469–81.

60. Rose L, Gönczy P. Polarity establishment, asymmetric division and segregation of fate determinants in early C. elegans embryos. Developmental Cell. WormBook; 2005;22:788–98.

61. Srinivasan DG, Fisk RM, Xu H, van den Heuvel S. A complex of LIN-5 and GPR proteins regulates G protein signaling and spindle function in C elegans. Gene Dev. Cold Spring Harbor Lab; 2003;17:1225–39.

62. Nguyen-Ngoc T, Afshar K, Gönczy P. Coupling of cortical dynein and G [alpha] proteins mediates spindle positioning in Caenorhabditis elegans. Nature cell biology. 2007.

63. Schierenberg E. Reversal of cellular polarity and early cell-cell interaction in the embryo of Caenorhabditis elegans. Dev Biol. 1987;122:452–63.

64. Cheng NN, Kirby CM, Kemphues KJ. Control of cleavage spindle orientation in Caenorhabditis elegans: the role of the genes par-2 and par-3. Genetics. 1995;139:549–59.

65. Whittle CM, McClinic KN, Ercan S, Zhang X, Green RD, Kelly WG, et al. The Genomic Distribution and Function of Histone Variant HTZ-1 during C. elegans Embryogenesis. Akhtar A, editor. PLoS Genet. Public Library of Science; 2008;4:e1000187.

66. Welch D, Meselson M. Evidence for the evolution of bdelloid rotifers without sexual reproduction or genetic exchange. Science. 2000;288:1211–5.

67. Butlin R. Evolution of sex: The costs and benefits of sex: new insights from old asexual lineages. Nature Reviews Genetics. 2002.

68. Blanc-Mathieu R, Perfus-Barbeoch L, Aury J-M, Da Rocha M, Gouzy J, Sallet E, et al. Hybridization and polyploidy enable genomic plasticity without sex in the most devastating plant-parasitic nematodes. Gojobori T, editor. PLoS Genet. 2017;13:e1006777–36.

69. Kimble J, Crittenden SL. Controls of Germline Stem Cells, Entry into Meiosis, and the Sperm/Oocyte Decision in Caenorhabditis elegans. http://dx.doi.org/10.1146/annurev.cellbio.23.090506.123326. Annual Reviews; 2007;23:405–33.

70. Lahl V, Halama C, Schierenberg E. Comparative and experimental embryogenesis of Plectidae (Nematoda). Development Genes and Evolution. 2003;213:18–27.

71. Brown HW, Cort WW. The Egg Production of Ascaris lumbricoides. The Journal of Parasitology. 1927;14:88.

72. HECHLER HC. Reproduction, Chromosome Number and Postembryonic Development of Panagrellus redivivus (Nematoda: Cephalobidae). 2001;:1–7.

73. Viera A, Page J, Rufas JS. Inverted Meiosis: The True Bugs as a Model to Study. Meiosis. Basel: Karger Publishers; 2009;5:137–56.

74. Beatty A, Morton D, Kemphues K. The C. elegans homolog of Drosophila Lethal giant larvae functions redundantly with PAR-2 to maintain polarity in the early embryo. Development. Oxford University Press for The Company of Biologists Limited; 2010;137:3995–4004.

75. Wirtz-Peitz F, Knoblich JA. Lethal giant larvae take on a life of their own. Trends in Cell Biology. 2006;16:234–41.

76. Knoblich JA. Mechanisms of asymmetric stem cell division. Cell. 2008;132:583–97.

77. Roegiers F, Jan YN. Asymmetric cell division. Current Opinion in Cell Biology. 2004;16:195–205.

78. Hyman AA. Centrosome movement in the early divisions of Caenorhabditis elegans: a cortical site determining centrosome position. J. Cell Biol. The Rockefeller University Press; 1989;109:1185–93.

79. Hyman AA, White JG. Determination of cell division axes in the early embryogenesis of Caenorhabditis elegans. J. Cell Biol. The Rockefeller University Press; 1987;105:2123–35.

80. Schulze J, Schierenberg E. Cellular pattern formation, establishment of polarity and segregation of colored cytoplasm in embryos of the nematode Romanomermis culicivorax. Dev Biol. 2008;315:426–36.

81. Brauchle M, Kiontke K, MacMenamin P, Fitch DHA, Piano F. Evolution of early embryogenesis in rhabditid nematodes. Dev Biol. 2009;335:253–62.

82. Tipney H, Hunter L. An introduction to effective use of enrichment analysis software. Hum. Genomics. BioMed Central; 2010;4:202–6.

83. Huang DW, Sherman BT, Lempicki RA. Bioinformatics enrichment tools: paths toward the comprehensive functional analysis of large gene lists. Nucleic Acids Res. 2009;37:1–13.

84. Lynch M. Destabilizing hybridization, general-purpose genotypes and geographic parthenogenesis. Q Rev Biol. 1984.

85. Lunt DH. Genetic tests of ancient asexuality in Root Knot Nematodes reveal recent hybrid origins. BMC Evol Biol. 2008;8:194–16.

86. Zhou T, Gu W, Wilke CO. Detecting positive and purifying selection at synonymous sites in yeast and worm. Mol Biol Evol. 2010;27:1912–22.

87. The complex hybrid origins of the root knot nematodes revealed through comparative genomics. 1st ed. PeerJ Inc; 2014;2:e356–25.

88. Szitenberg A, Salazar-Jaramillo L, Blok VC. Comparative genomics of apomictic root-knot nematodes: hybridization, ploidy, and dynamic genome change. bioRxiv. 2017.

89. Schiffer PH, Danchin E, Burnell AM, Schiffer A-M, Creevey C, Wong S, et al. Signatures of the evolution of parthenogenesis and cryptobiosis in the genomes of panagrolaimid nematodes. 2017.

90. Nicholas WL. The Biology of Free-Living Nematodes. Oxford University Press; 1984.

91. Nigon V. Developpement et reproduction des nematodes. In Traite de Zoologie, Tome IV (1965). 1965.

92. Schubert I. Chromosome evolution. Current Opinion in Plant Biology. 2007;10:109–15.

93. Wienberg J. The evolution of eutherian chromosomes. Current Opinion in Genetics & Development. 2004;14:657–66.

94. Félix M-A, Braendle C, Cutter AD. A Streamlined System for Species Diagnosis in Caenorhabditis (Nematoda: Rhabditidae) with Name Designations for 15 Distinct Biological Species. Goldstein B, editor. PLoS ONE. Public Library of Science; 2014;9:e94723.

95. Schierenberg E, Schulze J, Fusco G. Many roads lead to Rome: different ways to construct a nematode. Minelli A, editor. Key Themes in Evolutionary Developmental Biology. Cambridge: Cambridge University Press; 2008. pp. 261–80.

96. Schulze J, Schierenberg E. Evolution of embryonic development in nematodes. Evodevo. 2011;2:18.

97. Tintori SC, Osborne Nishimura E, Golden P, Lieb JD, Goldstein B. A Transcriptional Lineage of the Early C. elegans Embryo. Developmental Cell. 2016;38:430–44.

